# Virological characteristics of the SARS-CoV-2 JN.1 variant

**DOI:** 10.1101/2023.12.08.570782

**Authors:** Yu Kaku, Kaho Okumura, Miguel Padilla-Blanco, Yusuke Kosugi, Keiya Uriu, Alfredo A Hinay, Luo Chen, Arnon Plianchaisuk, Kouji Kobiyama, Ken J Ishii, The Genotype to Phenotype Japan (G2P-Japan) Consortium, Jiri Zahradnik, Jumpei Ito, Kei Sato

**Affiliations:** Division of Systems Virology, Department of Microbiology and Immunology, The Institute of Medical Science, The University of Tokyo, Tokyo, Japan; Faculty of Liberal Arts, Sophia University, Tokyo, Japan; First Medical Faculty at Biocev, Charles University, Vestec-Prague, Czechia; Departamento de Farmacia, Facultad de Ciencias de la Salud, Universidad Cardenal Herrera-CEU (UCH-CEU), CEU Universities, Valencia, Spain; Graduate School of Medicine, The University of Tokyo, Tokyo, Japan; Graduate School of Frontier Sciences, The University of Tokyo, Kashiwa, Japan; Division of Vaccine Science, Department of Microbiology and Immunology, The Institute of Medical Science, The University of Tokyo, Tokyo, Japan; International Vaccine Design Center, The Institute of Medical Science, The University of Tokyo, Tokyo, Japan; International Research Center for Infectious Diseases, The Institute of Medical Science, The University of Tokyo, Tokyo, Japan; Collaboration Unit for Infection, Joint Research Center for Human Retrovirus infection, Kumamoto University, Kumamoto, Japan; CREST, Japan Science and Technology Agency, Kawaguchi, Japan

## Abstract

The SARS-CoV-2 BA.2.86 lineage, first identified in August 2023, is phylogenetically distinct from the currently circulating SARS-CoV-2 Omicron XBB lineages, including EG.5.1 and HK.3. Comparing to XBB and BA.2, BA.2.86 carries more than 30 mutations in the spike (S) protein, indicating a high potential for immune evasion. BA.2.86 has evolved and its descendant, JN.1 (BA.2.86.1.1), emerged in late 2023. JN.1 harbors S:L455S and three mutations in non-S proteins. S:L455S is a hallmark mutation of JN.1: we have recently shown that HK.3 and other “FLip” variants carry S:L455F, which contributes to increased transmissibility and immune escape ability compared to the parental EG.5.1 variant. Here, we investigated the virological properties of JN.1.

## Text

The SARS-CoV-2 BA.2.86 lineage, first identified in August 2023, is phylogenetically distinct from the currently circulating SARS-CoV-2 Omicron XBB lineages, including EG.5.1 and HK.3. Comparing to XBB and BA.2, BA.2.86 carries more than 30 mutations in the spike (S) protein, indicating a high potential for immune evasion.^1-4^ BA.2.86 has evolved and its descendant, JN.1 (BA.2.86.1.1), emerged in late 2023. JN.1 harbors S:L455S and three mutations in non-S proteins (**Figure 1A**). S:L455S is a hallmark mutation of JN.1: we have recently shown that HK.3 and other “FLip” variants carry S:L455F, which contributes to increased transmissibility and immune escape ability compared to the parental EG.5.1 variant.^5^ Here, we investigated the virological properties of JN.1. We estimated the relative effective reproductive number (Re) of JN.1 using genomic surveillance data from France, the United Kingdom and Spain, where >25 sequences of JN.1 have been reported, using a Bayesian multinomial logistic model (**Figures 1B, 1C, Table S3**).^6^ The Re of JN.1 in these three countries was higher than that of BA.2.86.1 and HK.3, one of the XBB lineages with the highest growth advantage at the end of November 2023 (**Figure 1B**).^5^ These results suggest that JN.1 may soon become the dominant lineage worldwide. Indeed, by the end of November 2023, JN.1 has already overtaken HK.3 in France and Spain (**Figure 1C**).

**Figure 1.**
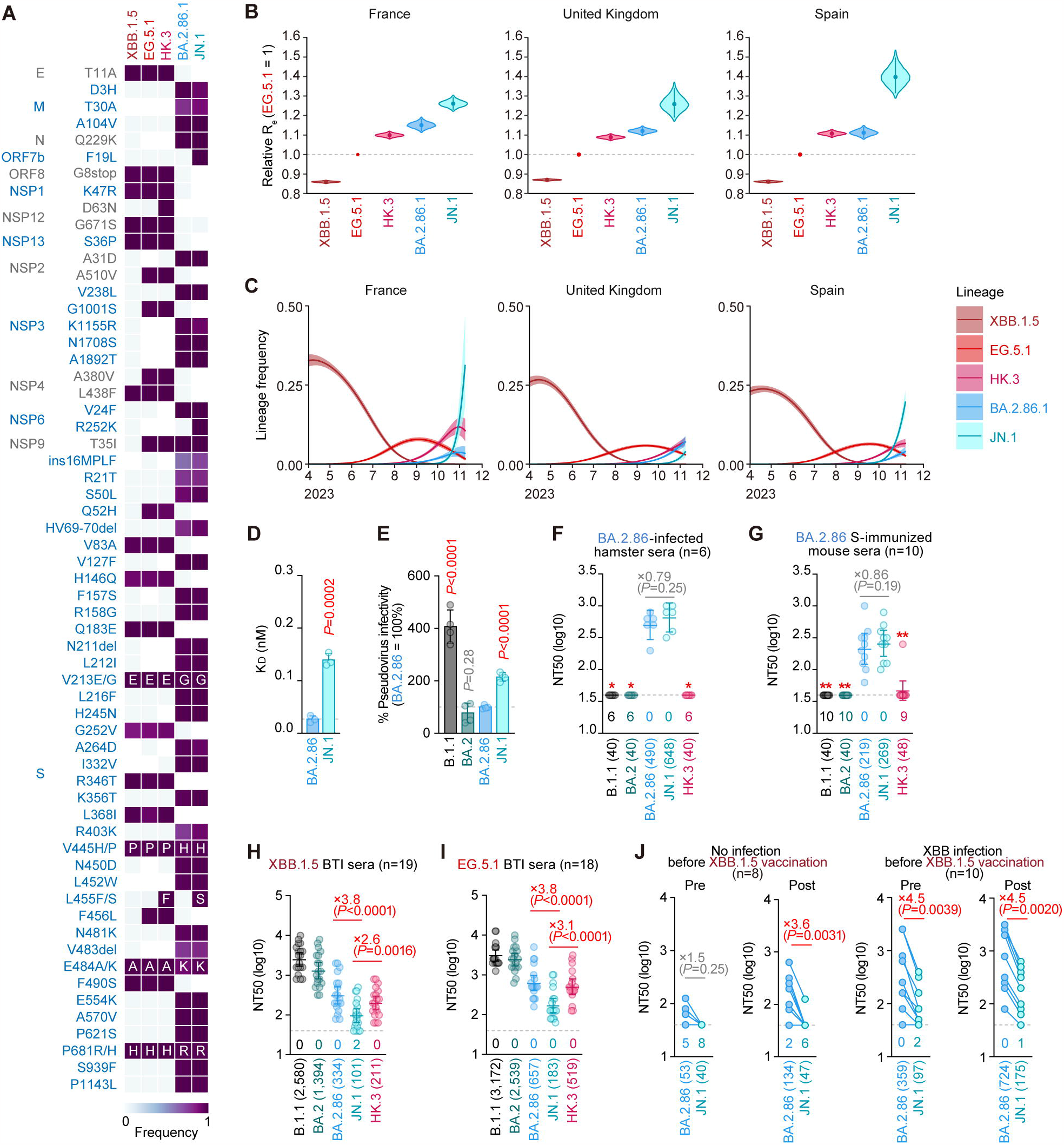
Virological features of JN.1. (**A**) Frequency of mutations in JN.1 and other lineages of interest. Only mutations with a frequency >0.5 in at least one but not all the representative lineages are shown. (**B**) Estimated relative Re of the variants of interest in France, United Kingdom, and Spain. The relative Re of EG.5.1 is set to 1 (horizontal dashed line). Violin, posterior distribution; dot, posterior mean; line, 95% Bayesian confidence interval. (**C**) Estimated epidemic dynamics of the variants of interest in France, United Kingdom, and Spain from April 1, 2023 to November 16, 2023. Countries are ordered according to the number of detected sequences of JN.1 from high to low. Line, posterior mean, ribbon, 95% Bayesian confidence interval. (**D**) Yeast surface display affinity between the RBD of the BA.2.86 SARS-CoV-2 variant or BA.2.86 that contained the L455S mutation and mACE2 was measured by yeast surface display. The dissociation constant (KD) value indicates the binding affinity of the RBD of the SARS-CoV-2 S protein to soluble ACE2 when expressed on yeast. Statistically significant differences versus BA.2.86 is determined by two-sided Student’s *t* tests. (**E**) Lentivirus-based pseudovirus assay. HOS-ACE2/TMPRSS2 cells were infected with pseudoviruses bearing each S protein of B.1.1 or BA.2 sublineages. The amount of input virus was normalized to the amount of HIV-1 p24 capsid protein. The percentage infectivity of B.1.1, BA.2 and JN.1 are compared to that of BA.2.86. The horizontal dash line indicates the mean value of the percentage infectivity of BA.2.86. Assays were performed in quadruplicate, and a representative result of four independent assays is shown. The presented data are expressed as the average ± SD. Each dot indicates the result of an individual replicate. Statistically significant differences versus BA.2.86 is determined by two-sided Student’s *t* tests. (**F**-**J**) Neutralization assay. Assays were performed with pseudoviruses harboring the S proteins of B.1.1, BA.2, BA.2.86, JN.1 and HK.3. The following sera were used: sera from six hamsters infected with BA.2.86 (**F**); sera from ten mice immunized with SARS-CoV-2 BA.2.86 S (**G**); convalescent sera from fully vaccinated individuals who had been infected with XBB.1.5 (eight 3-dose vaccinated donors, six 4-dose vaccinated donors, four 5-dose vaccinated donors and one 6-dose vaccinated donor. 19 donors in total) (**H**); and EG.5.1 (one 2-dose vaccinated donor, four 3-dose vaccinated donors, five 4-dose vaccinated donors, four 5-dose vaccinated donors and four 6-dose vaccinated donors. 18 donors in total) (**I**). Assays were also performed with pseudoviruses harboring the S proteins of BA.2.86 and JN.1. The following two sera were used: vaccinated sera from fully vaccinated individuals who had not been infected (8 donors) and vaccinated sera from fully vaccinated individuals who had been infected with XBB subvariants (after June, 2023) (10 donors). Sera were collected before vaccination (‘Pre’) and 3-4 weeks after XBB.1.5 monovalent vaccination (‘Post’) (**J**). Assays for each serum sample were performed in triplicate to determine the 50% neutralization titer (NT50).

Each dot represents one NT50 value, and the geometric mean and 95% confidence interval are shown. The number in parenthesis indicates the geometric mean of NT50 values. The horizontal dash line indicates the detection limit (40-fold) and the number of samples with neutralization titer under the limit are shown below the dash line. In **F**-**J**, statistically significant differences versus JN.1 were determined by two-sided Wilcoxon signed-rank tests, and p values are indicated in parentheses. The fold changes of NT50 from that of JN.1 are indicated with “X”. In **F** and **G**, *, p<0.05; **, p<0.01 versus JN.1.

The *in vitro* ACE2 binding assay^7^ showed that the dissociation constant (KD) value of the JN.1 receptor-binding domain (RBD) is significantly higher than that of the BA.2.86 RBD (**Figure 1D**), suggesting that S:L455S decreases the binding affinity to the human ACE2 receptor. In contrast, the pseudovirus assay showed that the infectivity of JN.1 is significantly higher than that of BA.2.86 (**Figure 1E**). This discrepancy (**Figures 1D, 1E**) would be due to the difference between monomeric RBD and trimerized whole S protein (see also **Supplementary Discussion**). We then performed a neutralization assay using rodent sera infected with BA.2.86 or immunized with BA.2.86 S protein. In both cases, the 50% neutralization titer (NT50) against JN.1 was comparable to that against BA.2.86 (**Figures 1F, 1G**), suggesting that S:L455S does not affect the antigenicity of BA.2.86. On the other hand, the NT50 of breakthrough infection (BTI) sera with XBB.1.5 and EG.5.1 against JN.1 was significantly lower than that of HK.3 (2.6-to 3.1-fold) and BA.2.86 (3.8-fold) (**Figures 1H, 1I**). Furthermore, JN.1 shows robust resistance to monovalent XBB.1.5 vaccine sera compared to BA.2.86 (**Figure 1J**). Taken together, these results suggest that JN.1 is one of the most immune-evading variants to date. Our results suggest that S:L455S contributes to increased immune evasion, which partly explains the increased Re of JN.1.

## Grants

Supported in part by AMED SCARDA Japan Initiative for World-leading Vaccine Research and Development Centers “UTOPIA” (JP223fa627001, to Ken J Ishii and Kei Sato), AMED SCARDA Program on R&D of new generation vaccine including new modality application (JP223fa727002, to Ken J Ishii and Kei Sato); AMED Research Program on Emerging and Re-emerging Infectious Diseases (JP22fk0108146, to Kei Sato; JP21fk0108494 to G2P-Japan Consortium and Kei Sato; JP21fk0108425, to Kei Sato; JP21fk0108432, to Kei Sato; JP22fk0108511, to G2P-Japan Consortium and Kei Sato; JP22fk0108516, to Kei Sato; JP22fk0108506, to Kei Sato); AMED Research Program on HIV/AIDS (JP22fk0410039, to Kei Sato); JST PRESTO (JPMJPR22R1, to Jumpei Ito); JST CREST (JPMJCR20H4, to Kei Sato); JSPS KAKENHI Fund for the Promotion of Joint International Research (International Leading Research) (JP23K20041, to G2P-Japan and Kei Sato); JSPS KAKENHI Grant-in-Aid for Early-Career Scientists (23K14526, to Jumpei Ito); JSPS Core-to-Core Program (A. Advanced Research Networks) (JPJSCCA20190008, Kei Sato); JSPS Research Fellow DC2 (22J11578, to Keiya Uriu); JSPS Research Fellow DC1 (23KJ0710, to Yusuke Kosugi); The Tokyo Biochemical Research Foundation (to Kei Sato); The Mitsubishi Foundation (to Kei Sato); International Joint Research Project of the Institute of Medical Science, the University of Tokyo (to Jiri Zahradnik); and the project of National Institute of Virology and Bacteriology, Programme EXCELES, funded by the European Union, Next Generation EU (LX22NPO5103, to Jiri Zahradnik). We wish to express our gratitude to CEU Universities and Santander Bank (Ayudas a la movilidad internacional de los investigadores en formación de la CEINDO) as well as to the Federation of European Biochemical Societies (FEBS; Short-Term Fellowship), for their financial support to Miguel Padilla-Blanco during the first and second part of his internship period at BIOCEV, respectively.

