## Supplementary Appendix for "Virological characteristics of the SARS-CoV-2 JN.1 variant"

**Table of Contents**

**Contents** Page


| | |
|---|---|
| **Supplementary Discussion** | 2 |
| **Materials and Methods** | 3-7 |
| Ethics statement | |
| Human serum collection | |
| Hamster serum collection | |
| Mouse serum collection | |
| Epidemic dynamics analysis and mutation frequency calculation | |
| Yeast surface display analysis | |
| Plasmid construction | |
| Cell culture | |
| Pseudovirus preparation | |
| Neutralization assay | |
| Data availability | |
| **Table S1.** Human infection sera used in this study | 8 |
| **Table S2.** Human XBB.1.5 vaccine sera used in this study | 9 |
| **Table S3.** Estimated relative Re and epidemic dynamics modeling parameters of the representative SARS-CoV-2 Omicron sublineages spreading in France, United Kingdom, and Spain from April 1, 2023 to November 16, 2023 | 10-15 |
| **Table S4.** Primers used in this study | 16 |
| **Figure S1.** Virological features of JN.1 | 17-18 |
| **Consortia** | 19 |
| **Acknowledgments** | 20 |
| **Supplemental References** | 21 |

### Supplementary Discussion

Infection receptor affinity is an extremely important factor in determining viral infectivity, but it is influenced by many factors. Lower infectivity for higher affinity mutations has been observed in our previous data[1] as well as in the work of others.[2] In addition, Han et al. have shown that the ACE2 binding affinity of the S RBD of the Beta variant is comparable to that of the Gamma variant, but the pseudovirus infectivity of the Gamma variant is lower than that of the Beta variant.[3] Similar to these previous studies, here we observed a discrepancy between the result of binding affinity of S RBD to the ACE2 receptor (**Figure S1D**) and that of pseudovirus infection (**Figure S1E**). We can speculate that spike stability, trimer packing and spike dynamics contribute to the observed phenomenon. Disentangling these effects is difficult and would be beyond the scope of this paper. We are working on this, but it will be the subject of future investigations. It seems that affinity plays a role mostly within a certain window of affinities. When the affinity is low, which is the case here, other factors may become important. The independent limitation, that should be mentioned, is the experimental approach, which is likely to be less stringent on affinity than in real-world transfer, where lower concentrations are present.

### Materials and Methods

#### Ethics statement

All protocols involving specimens from human subjects recruited at The Institute of Medical Science, Kyoto University, Interpark Kuramochi Clinic, Namikibashi Clinic and Wakaba Clinic was reviewed and approved by the Institutional Review Boards of The Institute of Medical Science (approval IDs: 2021-1-0416 and 2022-29-0915), Kyoto University (approval ID: G1309), Interpark Kuramochi Clinic (approval ID: G2021-004), Namikibashi Clinic and Wakaba Clinic (approval ID: 2022-29-0915), respectively. All human subjects provided written informed consent. All protocols for the use of human specimens were reviewed and approved by the Institutional Review Boards of The Institute of Medical Science, The University of Tokyo (approval IDs: 2021-1-0416, 2021-18-0617 and 2022-29-0915).

#### Human serum collection

Convalescent sera were collected from fully vaccinated individuals who had been infected with XBB.1.5 (eight 3-dose vaccinated, six 4-dose vaccinated, four 5-dose vaccinated and one 6-dose vaccinated; time interval between the last vaccination and infection, 44-524 days; 14-46 days after testing. n=19 in total; average age: 46.2 years, range: 15-74 years, 31.6% male) and EG.5.1 (one 2-dose vaccinated, four 3-dose vaccinated, five 4-dose vaccinated, four 5-dose vaccinated and four 6-dose vaccinated; time interval between the last vaccination and infection, 58–500 days; 4–27 days after testing. n=18 in total; average age: 55.1 years, range: 27–77 years, 50% male). XBB.1.5 monovalent vaccine sera from fully vaccinated individuals who had not been infected (eight donors. Average age, 64.4; range, 51–89; 37.5% male) and those from fully vaccinated individuals who had been infected with XBB subvariants (ten donors. Average age, 48.3; range, 32–61; 70% male) were collected before vaccination and three-four weeks (20–28 days) after vaccination. The SARS-CoV-2 variants were identified as previously described.[4-7] Sera were inactivated at 56°C for 30 minutes and stored at –80°C until use. The details of the convalescent sera are summarized in **Table S1 and Table S2**.

#### Hamster serum collection

Animal experiments were performed as previously described.[6-8] Briefly, 4-week male Syrian hamsters purchased from Japan SLC Inc. (Shizuoka, Japan) were inoculated with 10,000 50% tissue culture infectious dose ($TCID_{50}$) of SARS-CoV-2 BA.2.86 via the intranasal route under anesthesia. The anesthetics used for viral infection were injected into muscles as a mixture of 0.15 mg/kg medetomidine hydrochloride (Domitor®, Nippon Zenyaku Kogyo), 2.0 mg/kg midazolam (Dormicum®, Fujifilm Wako, Cat# 135-13791) and 2.5 mg/kg butorphanol (Vetorphale®, Meiji Seika Pharma) or 0.15 mg/kg medetomidine hydrochloride, 4.0 mg/kg alphaxaone (Alfaxan®, Jurox) and 2.5 mg/kg butorphanol at 16 days postinfection.

**Mouse serum collection**

The SARS-CoV-2 S-immunized mouse sera were prepared as previously described.[6,7] To prepare the immunogen, B16F10 cells (2,500,000 cells) were transfected with 5 µg S expression plasmid by PEI Max (Polysciences, Cat# 24765-1) according to the manufacturer's protocol. Two days posttransfection, the transfected cells were washed twice with PBS, and then the cell pellets were stored at –80°C (10,000,000 cells per stock). The expression of transfected S protein was verified by flow cytometry and western blot. BALB/c mice (female, 7 weeks old) were purchased from Japan SLC Inc. (Shizuoka, Japan). The mice were maintained under specific pathogen-free conditions. For the immunization, mice were subcutaneously immunized with the freeze-thawed S-expressing B16F10 cells in complete Freund's adjuvant (50%) (Sigma-Aldrich, Cat# F5881). Three weeks after immunization, blood was collected in BD Microtainer blood collection tubes (BD Biosciences, Cat# 365967) and sera were collected by centrifugation.

**Epidemic dynamics analysis and mutation frequency calculation**

In this study, we analyzed the viral genomic surveillance data deposited in the GISAID database (https://www.gisaid.org; downloaded on November 21, 2023). We used the data of SARS-CoV-2 collected from April 1, 2023 to November 16, 2023 in this analysis. We excluded the data of SARS-CoV-2 that i) lacks collection date and PANGO lineage information; ii) was retrieved from non-human animals; iii) was sampled by quarantine; iv) was sampled from the original passage; and v) whose genomic sequence is not longer than 28,000 base pairs and contains >2% of unknown (N) nucleotide sequences. In the downstream analysis, we only used sequences for PANGO lineages with >25 sequences in each country in the dataset. We modeled the epidemic dynamics of variants of interest in France, United Kingdom, and Spain, where >25 genomic sequences of JN.1 were detected. The daily frequency of each viral lineage was counted. Then, epidemic dynamics and $R_e$ value for each viral were subsequently estimated according to the Bayesian multinomial logistic model, described in our previous study.[7] Briefly, we estimated the logistic slope parameter $\beta_l$ for each lineage and then calculated a relative $R_e$ for each lineage ($r_l$) as $r_l = \exp(\gamma\beta_l)$ where $\gamma$ is the average viral generation time (2.1 days) (http://sonorouschocolate.com/covid19/index.php?title=Estimating_Generation_Time_Of_Omicron). For parameter estimation, the intercept and slope parameters of EG.5.1 were fixed at 0. The relative $R_e$ of EG.5.1 was fixed at 1, and that of other lineage was estimated with respect to that of EG.5.1. Parameter estimation was performed by using the Markov chain Monte Carlo (MCMC) approach implemented in CmdStan v2.33.1 (https://mc-stan.org) accessed through the CmdStanr v0.6.1 R interface (https://mc-stan.org/cmdstanr/). Four independent 5,000-step MCMC chains were run including 1,000-step warmup iterations. We confirmed that an estimated $\hat{R}$ convergence diagnostic value is <1.05 and bulk and tail effective sampling sizes are >200, indicating that all runs were successfully convergent. Information on the estimated parameters is summarized in **Table S3**. Mutation frequency of each lineage was calculated by dividing the number of sequences harboring the substitution of interest with the total number of sequences in each lineage.

#### Yeast surface display analysis

Yeast surface display was used to measure the interaction between the RBD of the SARS-CoV-2 BA.2.86 variant and JN.1 variant (RBD-BA.2.86 that contained the L455S mutation) and mACE2, following established protocols.[9-12] Site-directed mutagenesis by restriction-free cloning was conducted to incorporate the L455S mutation in the RBD-BA.2.86 (previously cloned in the pJYDC1 plasmid[1]), using the primers listed in **Table S4**. As previously outlined,[9-12] PCR reactions were performed using the KAPA HiFi HotStart ReadyMix kit (Roche, Cat# KK2601). The incorporation of the L455S mutations in the parental RBD-BA.2.86 was proved by Sanger sequencing, and the verified plasmid was then transformed into yeast Saccharomyces cerevisiae strain EBY100 (ATCC, MYA-4941) through electroporation and selection in SD-Trp plates. Yeast colonies were grown for 48 h in liquid culture (SDCAA, 30ºC, 220 rpm) and RBD expression in the yeast surface proceeded for 48h at 20ºC in 1/9 media. Yeasts expressing the RBD were washed with PBS supplemented with bovine serum albumin at a concentration of 1 g/L (PBSB). Subsequently, the cells were exposed to a range of mACE2 concentrations (2.5 pM to 5 nM, using two-fold serial dilutions) and 20 nM bilirubin (Sigma-Aldrich, Cat# 14370-1G), and the mixture was incubated for 30 min at 4ºC on darkness to promote the RBD-mACE2 binding. An additional washing with PBSB was performed prior to capturing the RBD expression and mACE2 signal using automated acquisition from a 96-well plate by the FACS CytoFLEX Flow Cytometer (Beckman Coulter). For the correct analysis, background binding signals were subtracted, and the fluorescence spill of eUnaG2 signals into the red channel was compensated. Finally, the data were fitted to a standard noncooperative Hill equation through nonlinear least-squares regression, utilizing Python v3.7 (https://www.python.org), as previously described.[9-12]

#### Plasmid construction

Plasmids expressing the SARS-CoV-2 spike proteins of B.1.1, BA.2, BA.2.86, and HK.3 and its derivative were prepared in our previous studies.[13-15] Plasmids expressing the spike protein of JN.1 was generated by site-directed overlap extension PCR using pC-SARS2-S BA.2.86 as the template and the primers listed in **Table S4**. The resulting PCR fragment was subcloned into the KpnI-NotI site of the pCAGGS vector[16] using In-Fusion HD Cloning Kit (Takara, Cat# Z9650N). Nucleotide sequences were determined by DNA sequencing services (Eurofins), and the sequence data were analyzed by SnapGene software v6.1.1 (www.snapgene.com).

#### Cell culture

The Lenti-X 293T cell line (Takara, Cat# 632180) and HOS-ACE2/TMPRSS2 cells (kindly provided by Dr. Kenzo Tokunaga), a derivative of HOS cells (a human osteosarcoma cell line; ATCC CRL-1543) stably expressing human ACE2 and TMPRSS2,[17,18] were maintained in Dulbecco's modified Eagle's medium (DMEM) (high glucose) (Wako, Cat# 044- 29765) containing

10% fetal bovine serum (Sigma-Aldrich Cat# 172012-500ML), 100 units penicillin and 100 ug/ml streptomycin (Sigma-Aldrich, Cat# P4333-100ML).

#### Pseudovirus preparation

Pseudoviruses were prepared as previously described.[13,15,19,20] Briefly, lentivirus (HIV-1)-based, luciferase-expressing reporter viruses were pseudotyped with the SARS-CoV-2 S. One prior day of transfection, the LentiX-293T cells were seeded at a density of 2 × $10^6$ cells. The LentiX-293T cells were cotransfected with 1 µg psPAX2-IN/HiBiT (a packaging plasmid encoding the HiBiT-tag-fused integrase[21]), 1 µg pWPI-Luc2 (a reporter plasmid encoding a firefly luciferase gene[21]) and 500 ng plasmids expressing parental S or its derivatives using TransIT-293 transfection reagent (Mirus, Cat# MIR2704) according to the manufacturer's protocol. Two days post transfection, the culture supernatants were harvested and filtrated. The amount of produced pseudovirus particles was quantified by the HiBiT assay using Nano Glo HiBiT lytic detection system (Promega, Cat# N3040) as previously described[21]. In this system, HiBiT peptide is produced with HIV-1 integrase and forms NanoLuc luciferase with LgBiT, which is supplemented with substrates. In each pseudovirus particle, the detected HiBiT value is correlated with the amount of the pseudovirus capsid protein, HIV-1 p24 protein.[21] Therefore, we calculated the amount of HIV-1 p24 capsid protein based on the HiBiT value measured, according to the previous paper.[21] To measure viral infectivity, the same amount of pseudovirus normalized with the HIV-1 p24 capsid protein was inoculated into HOS-ACE2/TMPRSS2 cells. At two days postinfection, the infected cells were lysed with a Bright-Glo luciferase assay system (Promega, Cat# E2620), and the luminescent signal produced by firefly luciferase reaction was measured using a GloMax explorer multimode microplate reader 3500 (Promega). The pseudoviruses were stored at –80°C until use.

#### Neutralization assay

Neutralization assays were performed as previously described.[10,11,13-15] The SARS-CoV-2 spike pseudoviruses (counting ~100,000 relative light units) were incubated with serially diluted (40-fold to 29,160-fold dilution at the final concentration) heat-inactivated sera at 37°C for 1 hour. Pseudoviruses without sera were included as controls. Then, 20 µl mixture of pseudovirus and serum was added to HOS-ACE2/TMPRSS2 cells (10,000 cells/100 µl) in a 96-well white plate. Two days post infection, the infected cells were lysed with a Bright-Glo luciferase assay system (Promega, Cat# E2620), and the luminescent signal was measured using a GloMax explorer multimode microplate reader 3500 (Promega). The assay of each serum sample was performed in triplicate, and the 50% neutralization titer ($NT_{50}$) was calculated using Prism 9 (GraphPad Software).

#### Data availability

The GISAID datasets used in this study are available from the GISAID database (https://www.gisaid.org; EPI_SET_231130we and EPI_SET_231130yx). The supplemental tables for the GISAID datasets are available in the GitHub repository (https://github.com/TheSatoLab/JN.1_short).

**Table S1. Human infection sera used in this study**

| SARS-CoV-2 infected | Donor ID | Sex | Age | Date of 1st vaccination (YYYY-MM-DD) | Date of 2nd vaccination (YYYY-MM-DD) | Date of 3rd vaccination (YYYY-MM-DD) | Date of 4th vaccination (YYYY-MM-DD) | Date of 5th vaccination (YYYY-MM-DD) | Date of 6th vaccination (YYYY-MM-DD) | Date of test (YYYY-MM-DD) | Date of sampling (YYYY-MM-DD) | Prior infection? |
|---|---|---|---|---|---|---|---|---|---|---|---|---|
| XBB.1.5 | 37306 | Female | 53 | 2021-04-27 (P) | 2021-05-18 (P) | 2022-02-01 (P) | 2022-07-30 (M) | 2022-12-17 (P) | | 2023-07-20 | 2023-08-08 | No |
| XBB.1.5 | 37097 | Female | 44 | NA (M) | 2021-08-16 (M) | 2022-05-13 (M) | | | | 2023-07-13 | 2023-08-11 | No |
| XBB.1.5 | 37598 | Female | 43 | 2021-04-28 (P) | 2021-05-19 (P) | 2022-11-08 (P) | 2022-07-08 (M) | 2022-12-27 (P) | | 2023-07-28 | 2023-08-11 | Yes |
| XBB.1.5 | 37072 | Female | 15 | 2021-09-25 (P) | 2021-10-18 (P) | 2022-05-02 (P) | | | | 2023-07-11 | 2023-08-11 | No |
| XBB.1.5 | 37071 | Female | 48 | 2021-10-01 (P) | 2021-11-01 (P) | 2022-05-06 (P) | | | | 2023-07-15 | 2023-08-11 | No |
| XBB.1.5 | 36845 | Male | 29 | 2021-09-01 (M) | 2021-09-29 (M) | 2022-05-27 (M) | | | | 2023-06-26 | 2023-08-11 | No |
| XBB.1.5 | 37229 | Female | 74 | 2021-06-24 (P) | 2021-07-15 (P) | 2022-02-16 (M) | 2022-07-20 (M) | 2023-03-25 (M) | | 2023-07-17 | 2023-08-01 | No |
| XBB.1.5 | 36708 | Male | 55 | 2021-08-07 (P) | 2021-08-27 (P) | 2022-04-14 (P) | | | | 2023-06-01 | 2023-07-09 | No |
| XBB.1.5 | 38084 | Male | 44 | 2021-09-13 (P) | 2021-10-05 (P) | 2022-07-29 (M) | | | | 2023-08-11 | 2023-09-02 | No |
| XBB.1.5 | 37998 | Female | 65 | 2021-08-04 (P) | 2021-08-30 (P) | 2022-03-19 (M) | 2022-09-02 (P) | 2022-12-24(PBA.4/5) | 2023-06-27 (P) | 2023-08-10 | 2023-09-02 | No |
| XBB.1.5 | 37798 | Female | 62 | 2021-03-17 (P) | 2021-04-09 (P) | 2021-12-23 (P) | 2022-07-28 (P) | 2023-06-17 (P) | | 2023-08-03 | 2023-08-20 | No |
| XBB.1.5 | 38061 | Female | 55 | 2021-08-17 (P) | 2021-09-18 (P) | 2022-04-02 (M) | 2022-10-14(PBA.1) | | | 2023-08-11 | 2023-09-04 | No |
| XBB.1.5 | 38019 | Female | 18 | 2021-09-07 (P) | 2021-10-07 (P) | 2022-04-28 (P) | 2022-12-27 (P) | | | 2023-08-10 | 2023-09-04 | No |
| XBB.1.5 | 38952 | Female | 54 | 2021-07-27 (M) | 2021-08-24 (M) | 2022-03-24 (P) | 2022-10-26 (P) | | | 2023-08-23 | 2023-09-10 | No |
| XBB.1.5 | 38871 | Male | 46 | 2021-09-12 (P) | 2021-10-03 (P) | 2022-04-08 (M) | 2022-11-05 (M) | | | 2023-08-22 | 2023-09-10 | No |
| XBB.1.5 | 38880 | Male | 51 | 2021-10-07 (P) | 2021-10-28 (P) | 2022-05-13 (M) | 2022-11-09 (PBA.4/5) | | | 2023-08-22 | 2023-09-16 | No |
| XBB.1.5 | 39018 | Female | 51 | 2021-07-29 (P) | 2021-08-23 (P) | 2022-03-19 (P) | | | | 2023-08-25 | 2023-09-13 | Yes |
| XBB.1.5 | 39019 | Female | 22 | 2021-07-26 (M) | 2021-08-23 (M) | 2022-03-19 (P) | | | | 2023-08-25 | 2023-09-13 | No |
| XBB.1.5 | 39296 | Male | 48 | 2021-07-15 (M) | 2021-08-23 (M) | 2022-03-24 (M) | 2022-12-25 (M) | | | 2023-09-01 | 2023-09-23 | No |
| EG.5 | P618 | Male | 54 | 2021-08-05 (P) | 2021-08-26 (P) | 2022-03-17 (P) | 2022-08-24 (P) | 2022-11-24 (P) | | 2023-08-23 | 2023-09-07 | No |
| EG.5 | KK-230801 | Male | 56 | 2021-07-16 (P) | 2021-08-06 (P) | 2022-03-03 (M) | 2022-08-09 (P) | | | 2023-07-28 | 2023-08-01 | No |
| EG.5 | 37301 | Female | 27 | 2021-08-01 (M) | 2022-03-06 (M) | | | | | 2023-07-19 | 2023-08-12 | No |
| EG.5 | 37330 | Female | 42 | 2021-09-25 (P) | 2021-10-16 (P) | 2022-05-13 (P) | | | | 2023-07-20 | 2023-08-13 | No |
| EG.5 | 38111 | Female | 50 | 2021-08-16 (P) | 2021-09-06 (P) | 2022-04-01 (P) | | | | 2023-08-12 | 2023-09-02 | No |
| EG.5 | 38197 | Female | 49 | 2021-08-20 (P) | 2021-09-10 (P) | 2022-03-25 (M) | 2022-11-18 (P) | | | 2023-08-13 | 2023-09-02 | No |
| EG.5 | 38217 | Male | 62 | 2021-07-04 (P) | 2021-07-25 (P) | 2022-02-26 (M) | 2022-08-06 (M) | 2022-11-20 (P) | | 2023-08-13 | 2023-09-02 | No |
| EG.5 | 37999 | Male | 67 | 2021-06-07 (P) | 2021-07-01 (P) | 2022-02-08 (P) | 2022-07-12 (P) | 2022-12-24 (PBA.4/5) | 2023-06-13 (MBA.4/5) | 2023-08-10 | 2023-09-02 | No |
| EG.5 | 38946 | Female | 65 | 2021-07-17 (P) | 2021-08-07 (P) | 2022-03-06 (M) | 2022-08-06 (M) | 2022-11-25 (P) | | 2023-08-23 | 2023-09-10 | No |
| EG.5 | 39025 | Male | 63 | 2021-08-01 (P) | 2021-08-22 (P) | 2022-03-07 (M) | 2022-08-10 (M) | 2022-11-26 (PBA.4/5) | 2023-06-06 (PBA.4/5) | 2023-08-26 | 2023-09-11 | No |
| EG.5 | 39288 | Male | 77 | 2021-06-09 (P) | 2021-07-01 (P) | 2022-02-05 (M) | 2022-07-12 (P) | 2022-11-18 (PBA.4/5) | 2023-05-30 (PBA.1) | 2023-08-31 | 2023-09-20 | No |
| EG.5 | 39301 | Male | 41 | 2021-08-26 (M) | 2021-09-23 (M) | 2022-04-24 (M) | 2022-10-09 (PBA.1) | | | 2023-09-01 | 2023-09-23 | No |
| EG.5 | 39314 | Female | 56 | 2021-07-25 (P) | 2021-08-22 (P) | 2022-03-03 (M) | 2022-08-06 (M) | 2022-11-06 (M) | 2023-04 (NA) | 2023-09-01 | 2023-09-21 | No |
| EG.5 | 39315 | Female | 60 | 2021-08-29 (P) | 2021-09-19 (P) | 2022-04-16 (M) | 2022-09-16 (P) | 2022-12-16 (PBA.4/5) | | 2023-08-31 | 2023-09-23 | No |
| EG.5 | 39316 | Male | 63 | 2021-07-31 (P) | 2021-08-21 (P) | 2022-04-23 (M) | 2022-09-30 (PBA.1) | | | 2023-08-31 | 2023-09-23 | No |
| EG.5 | 39330 | Male | 59 | 2021-08-24 (P) | 2021-09-14 (P) | 2022-03-26 (P) | 2022-10-23 (M) | | | 2023-08-31 | 2023-09-23 | No |
| EG.5 | 39503 | Female | 41 | 2021-08-24 (P) | 2021-09-15 (P) | 2022-04-27 (P) | | | | 2023-09-05 | 2023-09-25 | No |
| EG.5 | 39328 | Female | 59 | 2021-09-30 (P) | 2021-10-27 (P) | 2022-05-17 (P) | | | | 2023-08-31 | 2023-09-27 | Yes |

NA, not applicable.

P, Pfizer-BioNTech; M, Moderna

PBA.1, Pfizer/BioNTech bivalent vaccine - BA.1; PBA.4/5, Pfizer/BioNTech bivalent vaccine - BA.4-5; MBA.4/5, Moderna bivalent vaccine - BA.4-5

**Table S2. Human XBB.1.5 vaccine sera used in this study**

| Donor ID | Sex | Age | Date of 1st vaccination (YYYY-MM-DD) | Date of 2nd vaccination (YYYY-MM-DD) | Date of 3rd vaccination (YYYY-MM-DD) | Date of 4th vaccination (YYYY-MM-DD) | Date of 5th vaccination (YYYY-MM-DD) | Date of 6th vaccination (YYYY-MM-DD) | Date of sampling (before vaccination) (YYYY-MM-DD) | Date of XBB.1.5 vaccination (YYYY-MM-DD) | Date of sampling (after vaccination) (YYYY-MM-DD) | Time interval between vaccination and the second sampling | Prior infection? (YYYY-MM-DD) | Infected variant |
|---|---|---|---|---|---|---|---|---|---|---|---|---|---|---|
| 5165 | Female | 89 | 2021-05-29 (P) | 2021-06-21 (P) | 2022-02-16 (P) | 2022-07-17 (P) | 2022-11-27 (PBA4/5) | 2023-05-20 (PBA4/5) | 2023-09-29 | 2023-09-29 (XBB1.5) | 2023-10-26 | 27 | No | |
| 5166 | Male | 77 | 2021-06-05 (P) | 2021-07-04 (P) | 2022-03-05 (P) | 2022-08-06 (P) | 2022-11-20 (PBA4/5) | 2023-05-20 (PBA4/5) | 2023-09-29 | 2023-09-29 (XBB1.5) | 2023-10-21 | 22 | No | |
| 6783 | Male | 57 | 2021-06-23 (M) | 2021-07-21 (M) | 2022-02-11 (M) | 2022-10-15 (MBA1) | | | 2023-10-03 | 2023-10-03 (XBB1.5) | 2023-10-23 | 20 | No | |
| 6858 | Female | 62 | 2021-07-18 (P) | 2021-08-11 (P) | 2022-02-27 (M) | 2022-07-30 (P) | 2022-11-20 (PBA4/5) | | 2023-10-07 | 2023-10-07 (XBB1.5) | 2023-10-28 | 21 | No | |
| 192 | Female | 64 | 2021-04-22 (P) | 2021-05-13 (P) | 2022-01-15 (P) | 2022-07-16 (P) | 2022-11-24 (PBA4/5) | 2023-05-25 (PBA4/5) | 2023-10-02 | 2023-10-26 (PXBB1.5) | 2023-11-20 | 25 | No | |
| 1700 | Female | 53 | 2021-04-21 (P) | 2021-05-12 (P) | 2022-01-15 (P) | 2022-07-13 (P) | 2022-11-30 (PBA4/5) | 2023-06-23 (MBA4/5) | 2023-09-29 | 2023-10-18 (PXBB1.5) | 2023-11-13 | 26 | No | |
| 5555 | Female | 51 | 2021-07-14 (P) | 2021-08-14 (P) | 2022-02-22 (P) | 2022-07-23 (P) | 2022-12-03 (PBA4/5) | 2023-05-11 (PBA4/5) | 2023-09-30 | 2023-10-21 (PXBB1.5) | 2023-11-15 | 25 | No | |
| 5986 | Male | 62 | 2021-07-24 (P) | 2021-08-14 (P) | 2022-03-12 (M) | 2022-08-27 (M) | 2022-12-24 (PBA4/5) | 2023-06-03 (PBA4/5) | 2023-09-25 | 2023-10-21 (PXBB1.5) | 2023-11-13 | 23 | No | |
| KS | Male | 41 | 2021-06-17 (P) | 2021-07-07 (P) | 2022-03-28 (M) | 2022-10-27 (MBA.5) | | | 2023-09-19 | 2023-09-20 (XBB1.5) | 2023-10-14 | 24 | Yes (2023-06-29) | XBB.1.9 |
| KY | Female | 53 | 2021-08-18 (P) | 2021-09-08 (P) | 2022-04-13 (P) | 2022-10-21 (P) | | | 2023-09-25 | 2023-09-27 (XBB1.5) | 2023-10-20 | 23 | Yes (2023-07-24) | XBB.1.16 |
| KK | Male | 56 | 2021-07-16 (P) | 2021-08-06 (P) | 2022-03-03 (M) | 2022-08-09 (P) | | | 2023-09-25 | 2023-09-29 (XBB1.5) | 2023-10-24 | 25 | Yes (2023-07-17) | EG.5 |
| 2345 | Female | 52 | 2021-07-11 (P) | 2021-08-01 (P) | 2022-03-10 (P) | 2022-10-22 (PBA1) | | | 2023-10-06 | 2023-10-06 (XBB1.5) | 2023-10-27 | 21 | Yes (2023-07) | NA |
| 80 | Male | 61 | 2021-05-10 (P) | 2021-05-31 (P) | 2022-01-24 (M) | 2022-07-22 (BA4/5) | 2022-11-12 (BA4/5) | 2023-06-03 (BA4/5) | 2023-09-29 | 2023-09-30 (XBB1.5) | 2023-10-24 | 24 | Yes (2023-07) | NA |
| 90 | Male | 47 | 2021-05-11 (P) | 2021-06-02 (P) | 2022-01-25 (M) | 2022-08-02 (BA4/5) | 2022-11-09 (BA4/5) | | 2023-10-10 | 2023-10-11 (XBB1.5) | 2023-11-01 | 21 | Yes (2023-07) | NA |
| 100 | Male | 32 | 2021-05-18 (P) | 2021-06-08 (P) | 2022-01-31 (M) | 2022-07-17 (MBA4/5) | 2022-11-15 (PBA4/5) | 2023-05-27 (MBA4/5) | 2023-10-11 | 2023-10-14 (XBB1.5) | 2023-11-06 | 23 | Yes (2023-06) | NA |
| 286691 | Male | 49 | 2021-06-01(P) | 2021-06-22 (P) | 2022-03-21 (M) | | | | 2023-10-19 | 2023-10-19 (PXBB1.5) | 2023-11-16 | 28 | Yes (2023-08-22) | XBB.1.9 |
| 2439306 | Male | 43 | 2021-03-08 (P) | 2021-03-29 (P) | 2021-12-20 (P) | 2022-08-26 (M) | 2022-11-30 (M) | | 2023-10-23 | 2023-10-23 (PXBB1.5) | 2023-11-20 | 28 | Yes (2023-09-08) | NA |
| 4177 | Female | 49 | 2021-04-24 (P) | 2021-05-15 (P) | 2022-01-22 (P) | 2022-07-09 (P) | 2022-12-03 (PBA4/5) | 2023-06-13 (MBA4/5) | 2023-09-29 | 2023-10-14 (PXBB1.5) | 2023-11-10 | 27 | Yes (2023-08-29) | NA |

NA, not applicable.

P, Pfizer-BioNTech; M, Moderna

PBA.1, Pfizer/BioNTech bivalent vaccine - BA.1; PBA.4/5, Pfizer/BioNTech bivalent vaccine - BA.4-5; MBA.4/5, Moderna bivalent vaccine - BA.4-5

**Table S3. Estimated relative Re and epidemic dynamics modeling parameters of the representative SARS-CoV-2 Omicron sublineages spreading in France, United Kingdom, and Spain from April 1, 2023 to November 16, 2023**

| | | Relative $R_e$ (posterior values) | | | | | |
|---|---|---|---|---|---|---|---|
| PANGO lineage | Country | Mean | $2.5^{th}$ percentile | $97.5^{th}$ percentile | $\hat{R}$ | Bulk effective sample size | Tail effective sample size |
| JN.1 | France | 1.261 | 1.235 | 1.288 | 1.000 | 5346.8 | 9918.8 |
| BA.2.86.1 | France | 1.151 | 1.126 | 1.176 | 1.001 | 6214.5 | 10714.8 |
| JG.3 | France | 1.138 | 1.125 | 1.153 | 1.002 | 2344.1 | 7261.1 |
| JD.1.1.3 | France | 1.137 | 1.104 | 1.171 | 1.000 | 10574.6 | 11172.5 |
| HV.1 | France | 1.131 | 1.114 | 1.149 | 1.001 | 3801.7 | 8752.5 |
| FL.1.5.2 | France | 1.123 | 1.095 | 1.153 | 1.000 | 10187.7 | 11541.3 |
| HK.1 | France | 1.113 | 1.072 | 1.157 | 1.000 | 14479.9 | 11207.9 |
| HK.11 | France | 1.111 | 1.072 | 1.154 | 1.000 | 14040.8 | 11525.0 |
| JD.1.1 | France | 1.106 | 1.092 | 1.121 | 1.002 | 2532.5 | 6321.6 |
| HK.3 | France | 1.099 | 1.086 | 1.112 | 1.002 | 2019.8 | 6018.6 |
| EG.5.1.8 | France | 1.073 | 1.038 | 1.110 | 1.000 | 12571.8 | 11376.6 |
| JF.1 | France | 1.067 | 1.034 | 1.101 | 1.000 | 8141.2 | 10889.6 |
| XCH.1 | France | 1.056 | 1.041 | 1.071 | 1.002 | 2736.0 | 8239.2 |
| EG.6.1.1 | France | 1.052 | 1.018 | 1.087 | 1.000 | 10632.7 | 11268.8 |
| GK.1.1 | France | 1.046 | 1.027 | 1.066 | 1.001 | 5655.7 | 8565.8 |
| HK.6 | France | 1.039 | 1.021 | 1.058 | 1.001 | 4727.5 | 9063.8 |
| GS.4.1 | France | 1.034 | 1.027 | 1.042 | 1.008 | 666.2 | 2151.9 |
| XBB.1.16.17 | France | 1.031 | 1.016 | 1.045 | 1.002 | 2773.1 | 7045.1 |
| HF.1 | France | 1.024 | 1.001 | 1.049 | 1.001 | 7258.1 | 10434.1 |
| GK.4 | France | 1.023 | 0.996 | 1.052 | 1.000 | 10792.6 | 10993.5 |
| EG.5.1.6 | France | 1.022 | 1.010 | 1.035 | 1.002 | 1897.7 | 5082.1 |
| XBB.1.16.15 | France | 1.017 | 1.005 | 1.028 | 1.002 | 1700.7 | 5151.3 |
| DV.7.1.2 | France | 1.014 | 0.997 | 1.032 | 1.001 | 5193.8 | 8685.2 |
| FL.1.5.1 | France | 1.014 | 1.006 | 1.022 | 1.007 | 768.1 | 2764.6 |
| DV.7.1 | France | 1.013 | 1.002 | 1.025 | 1.004 | 1567.9 | 5709.6 |
| EG.10.1 | France | 1.008 | 0.994 | 1.023 | 1.002 | 2551.0 | 6627.0 |
| XBB.1.16.9 | France | 0.999 | 0.974 | 1.027 | 1.000 | 10345.6 | 9383.7 |
| EG.5.1.5 | France | 0.998 | 0.978 | 1.019 | 1.001 | 5986.3 | 9309.5 |
| EG.5.1.4 | France | 0.996 | 0.986 | 1.006 | 1.005 | 1193.7 | 3744.7 |
| EG.5.1.1 | France | 0.994 | 0.988 | 1.000 | 1.010 | 473.7 | 1393.2 |
| XBB.1.16.6 | France | 0.992 | 0.983 | 1.001 | 1.006 | 848.5 | 2827.2 |
| EG.5.1.3 | France | 0.992 | 0.986 | 0.997 | 1.012 | 375.4 | 1024.3 |
| FL.20 | France | 0.991 | 0.976 | 1.008 | 1.001 | 3257.0 | 8191.3 |
| XBB.2.3.11 | France | 0.991 | 0.983 | 0.999 | 1.007 | 730.5 | 2151.6 |
| XBB.1.16.11 | France | 0.988 | 0.980 | 0.995 | 1.007 | 620.7 | 1800.2 |
| EG.6.1 | France | 0.986 | 0.976 | 0.998 | 1.003 | 1533.2 | 5556.9 |
| GK.2.1 | France | 0.986 | 0.976 | 0.997 | 1.003 | 1474.2 | 4174.5 |
| JG.1 | France | 0.986 | 0.974 | 0.999 | 1.002 | 2321.3 | 6685.5 |
| GS.4 | France | 0.986 | 0.977 | 0.994 | 1.006 | 834.8 | 2164.3 |
| GK.1.3 | France | 0.985 | 0.970 | 1.000 | 1.002 | 3025.2 | 6657.0 |
| XBB.1.42.1 | France | 0.984 | 0.968 | 1.002 | 1.001 | 3419.9 | 8529.9 |
| GK.1 | France | 0.983 | 0.969 | 0.998 | 1.003 | 2488.4 | 6519.1 |
| GJ.1.2 | France | 0.979 | 0.970 | 0.988 | 1.006 | 902.2 | 2952.0 |
| GE.1 | France | 0.977 | 0.967 | 0.987 | 1.004 | 1271.7 | 3439.4 |
| GK.2 | France | 0.974 | 0.963 | 0.985 | 1.002 | 1612.8 | 5350.4 |
| XBB.1.5.70 | France | 0.972 | 0.955 | 0.991 | 1.001 | 4018.8 | 7579.4 |
| GS.5 | France | 0.967 | 0.949 | 0.988 | 1.002 | 4669.3 | 9049.7 |
| FW.1 | France | 0.967 | 0.951 | 0.984 | 1.001 | 3523.2 | 8826.9 |
| FL.15 | France | 0.965 | 0.954 | 0.976 | 1.004 | 1396.1 | 4303.1 |
| XBB.1.41.1 | France | 0.964 | 0.954 | 0.974 | 1.004 | 1233.7 | 4413.5 |
| XBB.1.16.14 | France | 0.959 | 0.947 | 0.971 | 1.003 | 1662.4 | 4891.7 |
| JA.1 | France | 0.956 | 0.942 | 0.971 | 1.002 | 2230.7 | 6972.2 |
| XBB.1.5.89 | France | 0.955 | 0.942 | 0.968 | 1.002 | 2045.8 | 6904.5 |
| HH.2 | France | 0.953 | 0.943 | 0.962 | 1.005 | 997.5 | 3481.7 |
| XBB.1.5.10 | France | 0.951 | 0.938 | 0.966 | 1.002 | 2656.9 | 7771.5 |
| FL.1.5 | France | 0.948 | 0.940 | 0.957 | 1.007 | 764.7 | 2559.1 |
| EG.2 | France | 0.945 | 0.932 | 0.958 | 1.002 | 2401.8 | 5830.6 |
| XBB.2.3.2 | France | 0.938 | 0.930 | 0.947 | 1.005 | 817.8 | 2777.5 |
| FU.1 | France | 0.932 | 0.922 | 0.941 | 1.004 | 1076.9 | 3106.2 |
| GJ.4 | France | 0.930 | 0.920 | 0.939 | 1.004 | 1128.9 | 3891.3 |
| XBB.1.16.2 | France | 0.929 | 0.916 | 0.943 | 1.003 | 1949.4 | 5842.0 |
| XBB.1.16.21 | France | 0.927 | 0.919 | 0.936 | 1.004 | 811.9 | 2914.3 |
| XBB.2.3.3 | France | 0.926 | 0.917 | 0.934 | 1.007 | 766.1 | 1968.2 |
| EG.4.5 | France | 0.924 | 0.911 | 0.937 | 1.002 | 1824.3 | 5973.3 |
| XBB.1.16 | France | 0.922 | 0.917 | 0.928 | 1.014 | 319.9 | 904.0 |
| XBB.1.16.19 | France | 0.922 | 0.914 | 0.930 | 1.006 | 785.8 | 2701.0 |

| | | | | | | | |
|---|---|---|---|---|---|---|---|
| XBB.2.3 | France | 0.917 | 0.910 | 0.924 | 1.009 | 507.0 | 1332.2 |
| XBB.1.5.59 | France | 0.915 | 0.906 | 0.923 | 1.007 | 743.1 | 2404.3 |
| FE.1.1 | France | 0.912 | 0.900 | 0.924 | 1.004 | 1597.4 | 4255.3 |
| FE.1.2 | France | 0.912 | 0.901 | 0.922 | 1.003 | 1333.6 | 4539.6 |
| GF.1 | France | 0.909 | 0.899 | 0.918 | 1.005 | 1044.1 | 3477.5 |
| FW.2 | France | 0.909 | 0.899 | 0.918 | 1.005 | 1002.9 | 3345.0 |
| XBB.1.16.1 | France | 0.908 | 0.901 | 0.914 | 1.010 | 453.6 | 1393.4 |
| EU.1.1 | France | 0.901 | 0.892 | 0.909 | 1.007 | 728.8 | 2502.2 |
| FY.6 | France | 0.898 | 0.889 | 0.907 | 1.005 | 880.8 | 3573.8 |
| FL.18 | France | 0.892 | 0.881 | 0.902 | 1.004 | 1224.4 | 4819.9 |
| BQ.1.1 | France | 0.890 | 0.877 | 0.901 | 1.004 | 1437.3 | 5640.0 |
| XBB.2.3.14 | France | 0.889 | 0.877 | 0.900 | 1.004 | 1430.0 | 5214.4 |
| EU.1.1.1 | France | 0.887 | 0.873 | 0.900 | 1.003 | 2063.5 | 7566.5 |
| XBB.1.5.37 | France | 0.883 | 0.876 | 0.890 | 1.009 | 534.9 | 1680.9 |
| XBB.1.5.4 | France | 0.883 | 0.871 | 0.895 | 1.002 | 1597.0 | 5689.9 |
| EG.1 | France | 0.881 | 0.875 | 0.887 | 1.010 | 447.3 | 1322.7 |
| XBB.1.5.1 | France | 0.878 | 0.866 | 0.889 | 1.002 | 1608.1 | 5350.4 |
| XBB.1.17.1 | France | 0.877 | 0.869 | 0.885 | 1.006 | 751.5 | 2657.9 |
| GB.1 | France | 0.876 | 0.870 | 0.883 | 1.009 | 513.4 | 1543.5 |
| EG.4 | France | 0.874 | 0.864 | 0.884 | 1.004 | 1202.1 | 3785.7 |
| EG.14 | France | 0.873 | 0.859 | 0.886 | 1.002 | 2387.9 | 8395.3 |
| FL.3.1 | France | 0.873 | 0.860 | 0.885 | 1.002 | 1934.8 | 5991.3 |
| FL.3 | France | 0.872 | 0.859 | 0.885 | 1.002 | 2245.7 | 6873.9 |
| FL.4 | France | 0.872 | 0.865 | 0.879 | 1.008 | 554.5 | 1785.6 |
| XBB.1.5.80 | France | 0.869 | 0.853 | 0.885 | 1.002 | 2917.0 | 7444.1 |
| XBB.1.9.1 | France | 0.869 | 0.863 | 0.875 | 1.012 | 407.1 | 1125.6 |
| FL.2 | France | 0.868 | 0.858 | 0.877 | 1.004 | 944.1 | 2817.0 |
| XBB.1.9.2 | France | 0.867 | 0.861 | 0.874 | 1.009 | 513.4 | 1469.2 |
| FY.1 | France | 0.865 | 0.847 | 0.882 | 1.001 | 3511.0 | 9408.5 |
| XBB.1.5.57 | France | 0.865 | 0.848 | 0.881 | 1.002 | 3912.7 | 8175.0 |
| FL.3.2 | France | 0.865 | 0.847 | 0.881 | 1.001 | 3548.7 | 9287.9 |
| FL.1 | France | 0.864 | 0.851 | 0.877 | 1.002 | 2127.7 | 5959.4 |
| XBB.1.22 | France | 0.863 | 0.843 | 0.882 | 1.001 | 5251.0 | 9305.6 |
| EG.1.3 | France | 0.861 | 0.850 | 0.871 | 1.003 | 1358.1 | 4653.1 |
| XBB.1.5 | France | 0.860 | 0.854 | 0.865 | 1.015 | 328.4 | 910.7 |
| XBB.1.5.48 | France | 0.860 | 0.845 | 0.873 | 1.002 | 2412.5 | 6417.0 |
| FL.5 | France | 0.857 | 0.848 | 0.866 | 1.005 | 902.1 | 3780.5 |
| XBB.1.5.66 | France | 0.857 | 0.835 | 0.876 | 1.002 | 5348.1 | 9743.4 |
| XBB.1.5.75 | France | 0.856 | 0.840 | 0.871 | 1.002 | 3203.2 | 8208.5 |
| GA.2 | France | 0.856 | 0.839 | 0.870 | 1.001 | 3107.6 | 7507.1 |
| EG.1.4 | France | 0.855 | 0.844 | 0.867 | 1.003 | 1662.8 | 6181.6 |
| FL.11 | France | 0.855 | 0.846 | 0.864 | 1.004 | 1105.5 | 3592.9 |
| FL.12 | France | 0.854 | 0.838 | 0.869 | 1.001 | 3216.9 | 8640.8 |
| FL.13 | France | 0.854 | 0.841 | 0.865 | 1.004 | 1876.5 | 5528.7 |
| FL.10 | France | 0.850 | 0.837 | 0.862 | 1.003 | 2000.5 | 5708.3 |
| XBB.1.5.94 | France | 0.850 | 0.838 | 0.860 | 1.004 | 1318.8 | 4381.8 |
| EG.13 | France | 0.849 | 0.840 | 0.859 | 1.004 | 1121.4 | 4863.4 |
| XBB.2.3.1 | France | 0.848 | 0.824 | 0.870 | 1.000 | 4891.7 | 11362.8 |
| XBB.1.5.15 | France | 0.846 | 0.833 | 0.858 | 1.002 | 2111.1 | 6246.4 |
| XBB.1.5.77 | France | 0.836 | 0.817 | 0.855 | 1.001 | 4418.9 | 8172.6 |
| XBB.1.5.24 | France | 0.836 | 0.819 | 0.853 | 1.002 | 3621.4 | 8866.5 |
| XBB.1.5.23 | France | 0.835 | 0.813 | 0.855 | 1.000 | 5560.0 | 8248.8 |
| CH.1.1 | France | 0.834 | 0.815 | 0.852 | 1.001 | 4426.7 | 8490.7 |
| XBB.1.5.65 | France | 0.833 | 0.811 | 0.854 | 1.000 | 6760.3 | 10615.4 |
| XBB.1.43.1 | France | 0.833 | 0.807 | 0.856 | 1.001 | 8042.7 | 11480.4 |
| XBB.1.5.13 | France | 0.830 | 0.820 | 0.841 | 1.003 | 1240.3 | 5331.9 |
| EK.4 | France | 0.827 | 0.798 | 0.854 | 1.000 | 10196.6 | 10551.1 |
| XBB.1.5.20 | France | 0.826 | 0.811 | 0.839 | 1.002 | 2661.5 | 6923.8 |
| XBB.1.5.90 | France | 0.824 | 0.793 | 0.853 | 1.000 | 11307.5 | 11185.5 |
| XBB.1.5.12 | France | 0.821 | 0.801 | 0.840 | 1.002 | 4484.2 | 9720.9 |
| XBB.1.5.7 | France | 0.820 | 0.806 | 0.833 | 1.001 | 2451.4 | 6419.2 |
| XBB.1.5.14 | France | 0.820 | 0.800 | 0.839 | 1.001 | 4863.6 | 10701.7 |
| EN.1.1 | France | 0.818 | 0.790 | 0.844 | 1.000 | 10272.4 | 9652.0 |
| XBB.1.5.100 | France | 0.817 | 0.788 | 0.844 | 1.001 | 10114.3 | 11001.5 |
| FL.27 | France | 0.816 | 0.795 | 0.836 | 1.002 | 5707.7 | 9895.5 |
| EN.1 | France | 0.810 | 0.783 | 0.835 | 1.000 | 10099.5 | 10910.0 |
| XBB.1.5.63 | France | 0.807 | 0.786 | 0.826 | 1.001 | 5712.9 | 11292.3 |
| XBB.1.5.21 | France | 0.789 | 0.757 | 0.819 | 1.001 | 14667.1 | 8793.8 |
| BQ.1.1.45 | France | 0.775 | 0.732 | 0.814 | 1.001 | 20306.8 | 10168.9 |
| JN.1 | United Kingdom | 1.261 | 1.200 | 1.327 | 1.000 | 12970.717 | 11622.4 |

| | | | | | | | |
|---|---|---|---|---|---|---|---|
| BA.2.86.2 | United Kingdom | 1.191 | 1.140 | 1.247 | 1.000 | 14149.1 | 10734.8 |
| JN.3 | United Kingdom | 1.165 | 1.134 | 1.198 | 1.000 | 13130.388 | 11550.0 |
| FL.15.1.1 | United Kingdom | 1.133 | 1.095 | 1.174 | 1.000 | 14548.8 | 11314.8 |
| BA.2.86.1 | United Kingdom | 1.121 | 1.105 | 1.137 | 1.000 | 4242.3 | 9853.2 |
| EG.5.1.8 | United Kingdom | 1.111 | 1.079 | 1.146 | 1.000 | 11746.6 | 10510.0 |
| JG.3 | United Kingdom | 1.111 | 1.095 | 1.126 | 1.000 | 4119.7 | 10504.9 |
| HV.1 | United Kingdom | 1.090 | 1.078 | 1.101 | 1.001 | 2420.6 | 6945.3 |
| HK.3 | United Kingdom | 1.089 | 1.077 | 1.100 | 1.001 | 2425.8 | 7483.9 |
| JD.1.1 | United Kingdom | 1.083 | 1.071 | 1.095 | 1.001 | 2852.5 | 7949.1 |
| JF.1 | United Kingdom | 1.059 | 1.046 | 1.073 | 1.001 | 3434.6 | 10492.1 |
| GS.4.1 | United Kingdom | 1.059 | 1.045 | 1.072 | 1.001 | 3216.6 | 9404.7 |
| FL.1.5.2 | United Kingdom | 1.058 | 1.044 | 1.074 | 1.001 | 5078.7 | 10761.6 |
| GA.4.1 | United Kingdom | 1.058 | 1.032 | 1.085 | 1.000 | 9976.5 | 10741.8 |
| JN.2 | United Kingdom | 1.057 | 1.041 | 1.075 | 1.000 | 5570.771 | 11344.9 |
| XCH.1 | United Kingdom | 1.045 | 1.029 | 1.061 | 1.000 | 4418.117 | 9818.2 |
| XBB.1.16.17 | United Kingdom | 1.041 | 1.023 | 1.060 | 1.000 | 6506.012 | 10219.3 |
| FY.5 | United Kingdom | 1.029 | 1.013 | 1.045 | 1.000 | 4837.0 | 8598.0 |
| EG.5.1.6 | United Kingdom | 1.029 | 1.020 | 1.037 | 1.003 | 1563.2 | 5255.3 |
| GS.4 | United Kingdom | 1.027 | 1.010 | 1.044 | 1.001 | 5226.6 | 9874.5 |
| HF.1 | United Kingdom | 1.025 | 1.015 | 1.036 | 1.002 | 2010.5 | 6498.3 |
| JM.2 | United Kingdom | 1.025 | 1.001 | 1.050 | 1.000 | 10220.5 | 10431.7 |
| DV.7.1 | United Kingdom | 1.024 | 1.013 | 1.035 | 1.001 | 2323.1 | 7650.4 |
| EG.10.1 | United Kingdom | 1.022 | 1.012 | 1.032 | 1.002 | 2180.6 | 6887.8 |
| GK.1.1 | United Kingdom | 1.022 | 1.008 | 1.036 | 1.001 | 3814.8 | 9389.5 |
| XBB.1.41.1 | United Kingdom | 1.022 | 1.008 | 1.035 | 1.001 | 3473.993 | 9216.1 |
| GW.5 | United Kingdom | 1.018 | 0.999 | 1.037 | 1.001 | 6768.5 | 9542.4 |
| FL.1.5.1 | United Kingdom | 1.012 | 1.006 | 1.019 | 1.004 | 731.7 | 2339.1 |
| XBB.1.16.15 | United Kingdom | 1.012 | 1.006 | 1.018 | 1.005 | 654.262 | 1892.0 |
| EG.5.1.3 | United Kingdom | 1.010 | 1.005 | 1.016 | 1.005 | 616.1 | 2218.4 |
| HW.1.1 | United Kingdom | 1.010 | 0.986 | 1.035 | 1.001 | 10980.2 | 10496.8 |
| XBB.1.16.11 | United Kingdom | 1.009 | 1.003 | 1.015 | 1.005 | 608.717 | 1793.0 |
| FL.20.1 | United Kingdom | 1.006 | 0.993 | 1.020 | 1.001 | 2964.1 | 8902.6 |
| JG.2 | United Kingdom | 1.005 | 0.984 | 1.028 | 1.000 | 6962.1 | 10304.4 |
| FL.24 | United Kingdom | 1.004 | 0.985 | 1.025 | 1.001 | 6189.6 | 10871.0 |
| EG.5.1.5 | United Kingdom | 1.004 | 0.986 | 1.023 | 1.000 | 6238.4 | 9982.2 |
| XBB.1.16.6 | United Kingdom | 1.004 | 0.997 | 1.010 | 1.004 | 722.224 | 2241.6 |
| EU.1.1.3 | United Kingdom | 1.004 | 0.985 | 1.024 | 1.001 | 5994.6 | 9457.4 |
| GK.1 | United Kingdom | 1.003 | 0.982 | 1.026 | 1.000 | 8353.2 | 10550.0 |
| HK.6 | United Kingdom | 1.002 | 0.995 | 1.009 | 1.003 | 897.2 | 3819.6 |
| EG.5.1.1 | United Kingdom | 1.001 | 0.996 | 1.006 | 1.006 | 476.9 | 1376.2 |
| EG.9.1 | United Kingdom | 1.001 | 0.979 | 1.023 | 1.001 | 7674.7 | 10485.7 |
| GJ.5.1 | United Kingdom | 0.999 | 0.984 | 1.015 | 1.001 | 4978.7 | 10232.0 |
| DV.7.1.2 | United Kingdom | 0.998 | 0.986 | 1.010 | 1.001 | 2927.0 | 8617.3 |
| HK.11 | United Kingdom | 0.997 | 0.985 | 1.010 | 1.001 | 2850.7 | 7246.2 |
| GK.2.1 | United Kingdom | 0.996 | 0.984 | 1.009 | 1.002 | 2807.5 | 8557.2 |
| EG.5.1.4 | United Kingdom | 0.996 | 0.989 | 1.003 | 1.004 | 896.0 | 3092.2 |
| EG.5.1.7 | United Kingdom | 0.995 | 0.982 | 1.009 | 1.001 | 3242.3 | 8895.1 |
| XCL | United Kingdom | 0.992 | 0.977 | 1.008 | 1.001 | 4078.656 | 8203.1 |
| GK.2 | United Kingdom | 0.992 | 0.977 | 1.007 | 1.001 | 4549.7 | 9592.2 |
| GE.1 | United Kingdom | 0.989 | 0.983 | 0.995 | 1.005 | 612.2 | 2001.0 |
| JM.1 | United Kingdom | 0.988 | 0.976 | 1.001 | 1.000 | 3449.9 | 8636.6 |
| XBB.1.42.1 | United Kingdom | 0.986 | 0.969 | 1.004 | 1.001 | 6056.527 | 8264.4 |
| FW.1.1 | United Kingdom | 0.984 | 0.970 | 0.999 | 1.001 | 4385.3 | 9270.6 |
| XBB.2.3.8 | United Kingdom | 0.983 | 0.967 | 0.999 | 1.001 | 5683.785 | 8935.2 |
| GJ.1.2 | United Kingdom | 0.982 | 0.974 | 0.990 | 1.003 | 1153.5 | 4680.6 |
| XBB.1.16.16 | United Kingdom | 0.981 | 0.963 | 1.000 | 1.000 | 7765.176 | 10266.3 |
| GE.1.3 | United Kingdom | 0.978 | 0.967 | 0.990 | 1.000 | 2803.0 | 7165.7 |
| GS.3 | United Kingdom | 0.976 | 0.965 | 0.988 | 1.001 | 2474.9 | 7975.7 |
| XBB.1.16.9 | United Kingdom | 0.976 | 0.966 | 0.987 | 1.001 | 2151.291 | 6371.0 |
| XBB.2.3.11 | United Kingdom | 0.970 | 0.963 | 0.978 | 1.003 | 1058.262 | 2945.9 |
| XBB.1.22.1 | United Kingdom | 0.965 | 0.954 | 0.976 | 1.001 | 2194.310 | 7518.9 |
| FL.20 | United Kingdom | 0.965 | 0.949 | 0.982 | 1.000 | 5330.5 | 10069.7 |
| EG.6.1 | United Kingdom | 0.965 | 0.958 | 0.972 | 1.004 | 889.8 | 3179.8 |
| XBB.1.16.14 | United Kingdom | 0.963 | 0.948 | 0.979 | 1.001 | 4301.349 | 8668.3 |
| FL.1.5 | United Kingdom | 0.963 | 0.952 | 0.974 | 1.001 | 2762.7 | 7134.1 |
| HK.9 | United Kingdom | 0.962 | 0.949 | 0.974 | 1.001 | 2945.7 | 8574.4 |
| XBB.1.16.8 | United Kingdom | 0.960 | 0.947 | 0.974 | 1.001 | 2957.535 | 8374.4 |
| XBB.2.3.3 | United Kingdom | 0.959 | 0.950 | 0.969 | 1.003 | 1772.741 | 6112.5 |
| FL.15 | United Kingdom | 0.955 | 0.942 | 0.969 | 1.001 | 3541.2 | 8701.4 |
| FY.1.2 | United Kingdom | 0.951 | 0.943 | 0.959 | 1.004 | 1188.0 | 3869.4 |

| | | | | | | | |
|---|---|---|---|---|---|---|---|
| FU.1 | United Kingdom | 0.950 | 0.943 | 0.958 | 1.003 | 982.7 | 3916.8 |
| EG.7 | United Kingdom | 0.949 | 0.940 | 0.958 | 1.002 | 1547.6 | 5253.1 |
| FY.4.1.2 | United Kingdom | 0.944 | 0.933 | 0.956 | 1.001 | 2920.2 | 7824.2 |
| JF.2 | United Kingdom | 0.944 | 0.932 | 0.956 | 1.001 | 2505.1 | 7185.4 |
| XBB.1.16.21 | United Kingdom | 0.943 | 0.936 | 0.950 | 1.004 | 844.435 | 3147.8 |
| XBB.1.16.19 | United Kingdom | 0.942 | 0.935 | 0.949 | 1.003 | 836.826 | 3137.9 |
| FU.2 | United Kingdom | 0.940 | 0.928 | 0.952 | 1.001 | 2327.7 | 7676.5 |
| EG.2 | United Kingdom | 0.939 | 0.930 | 0.949 | 1.002 | 1728.3 | 7316.2 |
| XBB.1.16 | United Kingdom | 0.939 | 0.935 | 0.944 | 1.010 | 316.805 | 689.5 |
| XBB.2.3.2 | United Kingdom | 0.937 | 0.929 | 0.945 | 1.003 | 1032.457 | 3460.2 |
| FE.1.2 | United Kingdom | 0.936 | 0.927 | 0.945 | 1.002 | 1509.2 | 6053.3 |
| XBB.1.5.28 | United Kingdom | 0.936 | 0.924 | 0.948 | 1.002 | 3331.605 | 8725.9 |
| HH.2 | United Kingdom | 0.936 | 0.923 | 0.948 | 1.001 | 3007.3 | 7751.8 |
| XBB.1.16.3 | United Kingdom | 0.936 | 0.926 | 0.945 | 1.001 | 1819.631 | 5508.2 |
| XBB.1.16.18 | United Kingdom | 0.934 | 0.924 | 0.944 | 1.002 | 1577.716 | 5965.4 |
| XBB.1.16.2 | United Kingdom | 0.933 | 0.925 | 0.940 | 1.003 | 917.726 | 2919.5 |
| XBB.1.16.1 | United Kingdom | 0.930 | 0.925 | 0.935 | 1.005 | 473.685 | 1249.1 |
| XBB.2.3 | United Kingdom | 0.930 | 0.924 | 0.936 | 1.005 | 573.097 | 1483.4 |
| EG.1.6 | United Kingdom | 0.927 | 0.915 | 0.939 | 1.001 | 2329.2 | 8647.2 |
| FL.4 | United Kingdom | 0.917 | 0.911 | 0.923 | 1.005 | 607.3 | 1805.9 |
| XBB.1.5.49 | United Kingdom | 0.914 | 0.903 | 0.924 | 1.001 | 2074.654 | 6515.6 |
| HZ.2 | United Kingdom | 0.911 | 0.900 | 0.923 | 1.002 | 2155.8 | 6933.2 |
| EG.1 | United Kingdom | 0.910 | 0.904 | 0.916 | 1.004 | 631.7 | 2000.4 |
| XBB.1.5.59 | United Kingdom | 0.910 | 0.902 | 0.917 | 1.002 | 1080.864 | 3346.1 |
| XBB.1.5.98 | United Kingdom | 0.905 | 0.892 | 0.917 | 1.001 | 3065.631 | 6807.0 |
| EG.13 | United Kingdom | 0.904 | 0.892 | 0.917 | 1.001 | 2597.7 | 8374.2 |
| EU.1.1 | United Kingdom | 0.903 | 0.893 | 0.912 | 1.001 | 1829.9 | 5967.2 |
| FL.13 | United Kingdom | 0.900 | 0.887 | 0.913 | 1.001 | 3261.1 | 9378.6 |
| EG.1.4 | United Kingdom | 0.900 | 0.887 | 0.912 | 1.002 | 2639.0 | 8316.5 |
| FL.2 | United Kingdom | 0.894 | 0.885 | 0.904 | 1.001 | 1836.0 | 6399.5 |
| XBB.1.17.1 | United Kingdom | 0.893 | 0.881 | 0.905 | 1.001 | 2912.088 | 7885.6 |
| EK.2.1 | United Kingdom | 0.892 | 0.881 | 0.902 | 1.001 | 1966.6 | 7647.3 |
| XBB.1.9.1 | United Kingdom | 0.890 | 0.885 | 0.896 | 1.005 | 515.794 | 1525.6 |
| FL.3.3 | United Kingdom | 0.888 | 0.879 | 0.896 | 1.002 | 1270.7 | 3734.4 |
| CH.1.1.1 | United Kingdom | 0.887 | 0.875 | 0.898 | 1.001 | 2324.5 | 7348.1 |
| FL.5 | United Kingdom | 0.885 | 0.875 | 0.895 | 1.001 | 2108.3 | 6347.3 |
| FZ.1.1 | United Kingdom | 0.885 | 0.875 | 0.895 | 1.002 | 1618.0 | 5810.2 |
| XBB.1.5.37 | United Kingdom | 0.885 | 0.876 | 0.893 | 1.002 | 1258.126 | 4618.1 |
| XBB.1.9.2 | United Kingdom | 0.884 | 0.878 | 0.890 | 1.004 | 674.714 | 2277.8 |
| XBB.1.5.15 | United Kingdom | 0.875 | 0.858 | 0.891 | 1.000 | 5060.058 | 10110.7 |
| DV.7 | United Kingdom | 0.873 | 0.859 | 0.886 | 1.001 | 3365.1 | 9546.1 |
| FL.3.2 | United Kingdom | 0.873 | 0.861 | 0.884 | 1.001 | 2470.7 | 7319.8 |
| XBB.1.5 | United Kingdom | 0.870 | 0.865 | 0.875 | 1.008 | 392.123 | 987.2 |
| XBB.1.5.1 | United Kingdom | 0.868 | 0.849 | 0.884 | 1.001 | 4675.291 | 10292.3 |
| FL.10 | United Kingdom | 0.867 | 0.852 | 0.880 | 1.001 | 3754.5 | 10104.0 |
| XBB.1.5.24 | United Kingdom | 0.866 | 0.852 | 0.879 | 1.001 | 3374.039 | 8606.9 |
| XBB.2.3.1 | United Kingdom | 0.865 | 0.848 | 0.881 | 1.001 | 5862.313 | 10002.1 |
| XBB.1.5.16 | United Kingdom | 0.864 | 0.848 | 0.879 | 1.000 | 5670.026 | 10199.5 |
| FL.18 | United Kingdom | 0.864 | 0.844 | 0.882 | 1.001 | 7092.1 | 11131.6 |
| XBB.1.5.4 | United Kingdom | 0.860 | 0.842 | 0.876 | 1.001 | 5277.884 | 10342.7 |
| FL.3.1 | United Kingdom | 0.859 | 0.850 | 0.868 | 1.001 | 1540.1 | 5144.0 |
| FL.3 | United Kingdom | 0.859 | 0.850 | 0.867 | 1.002 | 1308.8 | 4148.9 |
| XBB.1.5.46 | United Kingdom | 0.858 | 0.836 | 0.877 | 1.000 | 8083.794 | 10343.1 |
| XBB.1.5.47 | United Kingdom | 0.857 | 0.837 | 0.876 | 1.001 | 7176.860 | 10951.9 |
| XBB.1.5.20 | United Kingdom | 0.857 | 0.841 | 0.871 | 1.001 | 4019.232 | 8652.5 |
| XBB.1.5.12 | United Kingdom | 0.854 | 0.839 | 0.867 | 1.001 | 3540.532 | 8538.1 |
| EG.1.3 | United Kingdom | 0.853 | 0.840 | 0.865 | 1.001 | 3354.0 | 8731.6 |
| XBB.1.5.14 | United Kingdom | 0.852 | 0.831 | 0.872 | 1.000 | 8777.272 | 10335.9 |
| XBB.1.5.52 | United Kingdom | 0.850 | 0.835 | 0.865 | 1.000 | 5208.829 | 9333.5 |
| XBB.1.5.7 | United Kingdom | 0.849 | 0.840 | 0.858 | 1.002 | 1498.355 | 5465.8 |
| FL.9 | United Kingdom | 0.848 | 0.835 | 0.860 | 1.001 | 3258.7 | 7633.9 |
| EM.1 | United Kingdom | 0.846 | 0.830 | 0.862 | 1.000 | 5509.5 | 11004.9 |
| XBB.1.5.18 | United Kingdom | 0.840 | 0.829 | 0.852 | 1.001 | 2774.509 | 6909.0 |
| XBB.1.5.13 | United Kingdom | 0.825 | 0.812 | 0.838 | 1.000 | 3518.677 | 8581.2 |
| DV.6 | United Kingdom | 0.822 | 0.802 | 0.841 | 1.000 | 9132.1 | 10981.4 |
| XBB.1.5.48 | United Kingdom | 0.822 | 0.800 | 0.842 | 1.001 | 8855.344 | 10284.0 |
| XBB.1.5.97 | United Kingdom | 0.816 | 0.783 | 0.846 | 1.000 | 16913.706 | 12853.4 |
| CH.1.1 | United Kingdom | 0.780 | 0.745 | 0.812 | 1.001 | 15862.6 | 12718.2 |
| DV.1.1 | United Kingdom | 0.746 | 0.704 | 0.785 | 1.000 | 21569.4 | 13022.3 |
| JN.1 | Spain | 1.399 | 1.326 | 1.481 | 1.000 | 14401.668 | 10998.2 |

| | | | | | | | |
|---|---|---|---|---|---|---|---|
| JN.3 | Spain | 1.148 | 1.114 | 1.184 | 1.001 | 13499.329 | 11271.7 |
| HV.1 | Spain | 1.128 | 1.108 | 1.150 | 1.000 | 7069.042 | 10976.3 |
| JG.3 | Spain | 1.124 | 1.109 | 1.138 | 1.000 | 4002.763 | 7585.2 |
| BA.2.86.1 | Spain | 1.112 | 1.090 | 1.136 | 1.000 | 7059.561 | 11134.8 |
| HK.3 | Spain | 1.108 | 1.095 | 1.121 | 1.000 | 3020.590 | 8242.9 |
| JD.1.1 | Spain | 1.087 | 1.076 | 1.099 | 1.000 | 2532.721 | 6615.7 |
| DV.7.1.3 | Spain | 1.072 | 1.045 | 1.101 | 1.000 | 10607.001 | 10566.5 |
| XCH.1 | Spain | 1.055 | 1.040 | 1.071 | 1.000 | 4468.807 | 9749.0 |
| GS.4.1 | Spain | 1.051 | 1.031 | 1.071 | 1.000 | 7904.486 | 10602.5 |
| DV.7.1.1 | Spain | 1.041 | 1.019 | 1.064 | 1.000 | 8256.106 | 11294.3 |
| EG.5.1.6 | Spain | 1.035 | 1.020 | 1.050 | 1.000 | 3562.025 | 8266.3 |
| HK.6 | Spain | 1.030 | 1.019 | 1.041 | 1.001 | 2355.210 | 7571.0 |
| XBB.1.16.15 | Spain | 1.030 | 1.015 | 1.045 | 1.001 | 4503.115 | 10004.8 |
| EG.5.1.2 | Spain | 1.026 | 0.999 | 1.054 | 1.000 | 9953.186 | 11377.6 |
| JG.1 | Spain | 1.023 | 1.002 | 1.045 | 1.000 | 7579.159 | 11152.1 |
| EG.5.1.1 | Spain | 1.022 | 1.014 | 1.029 | 1.001 | 1114.614 | 3228.8 |
| EG.10.1 | Spain | 1.021 | 1.005 | 1.036 | 1.000 | 4727.737 | 10085.2 |
| FL.1.5.1 | Spain | 1.020 | 1.010 | 1.029 | 1.001 | 1754.113 | 4984.7 |
| XCL | Spain | 1.017 | 0.993 | 1.043 | 1.000 | 8725.796 | 10058.6 |
| EG.5.1.4 | Spain | 1.016 | 1.006 | 1.027 | 1.000 | 2596.898 | 7315.6 |
| EG.5.1.3 | Spain | 1.010 | 1.004 | 1.017 | 1.002 | 875.944 | 2600.1 |
| XBB.1.16.6 | Spain | 1.008 | 0.998 | 1.019 | 1.001 | 2041.079 | 6452.6 |
| XBB.1.42.1 | Spain | 1.003 | 0.982 | 1.026 | 1.000 | 7446.504 | 10769.7 |
| GK.2 | Spain | 0.996 | 0.981 | 1.012 | 1.000 | 3778.040 | 9498.2 |
| DV.7.1.2 | Spain | 0.992 | 0.979 | 1.006 | 1.000 | 3177.588 | 7955.7 |
| DV.7.1 | Spain | 0.991 | 0.984 | 0.997 | 1.002 | 755.657 | 2349.2 |
| GK.1 | Spain | 0.990 | 0.978 | 1.004 | 1.001 | 2756.163 | 5594.7 |
| GK.2.1 | Spain | 0.990 | 0.982 | 0.998 | 1.001 | 1306.669 | 3395.2 |
| GE.1 | Spain | 0.981 | 0.972 | 0.991 | 1.001 | 1677.225 | 5938.9 |
| XBB.1.16.11 | Spain | 0.977 | 0.970 | 0.983 | 1.002 | 785.475 | 2110.7 |
| GJ.1.2 | Spain | 0.977 | 0.963 | 0.990 | 1.001 | 3227.443 | 8443.6 |
| EG.6.1 | Spain | 0.974 | 0.963 | 0.985 | 1.001 | 2522.661 | 6631.3 |
| EG.5.1.5 | Spain | 0.972 | 0.962 | 0.983 | 1.001 | 1955.619 | 5410.1 |
| FL.15 | Spain | 0.972 | 0.957 | 0.988 | 1.001 | 4377.561 | 8966.1 |
| EG.11 | Spain | 0.971 | 0.954 | 0.990 | 1.000 | 5517.873 | 9842.6 |
| FL.20 | Spain | 0.969 | 0.962 | 0.977 | 1.001 | 1079.901 | 3215.9 |
| DV.7 | Spain | 0.962 | 0.945 | 0.981 | 1.000 | 6090.266 | 10579.0 |
| FL.1.5 | Spain | 0.958 | 0.948 | 0.968 | 1.001 | 1991.473 | 4290.5 |
| XBB.2.3.11 | Spain | 0.955 | 0.940 | 0.972 | 1.000 | 3878.243 | 8699.8 |
| FU.2 | Spain | 0.953 | 0.937 | 0.970 | 1.001 | 5003.119 | 9820.0 |
| GJ.2 | Spain | 0.953 | 0.939 | 0.967 | 1.001 | 2965.619 | 7732.5 |
| XBB.1.5.72 | Spain | 0.952 | 0.940 | 0.964 | 1.001 | 2187.429 | 7194.9 |
| FY.1.2 | Spain | 0.950 | 0.938 | 0.963 | 1.001 | 2909.558 | 7353.7 |
| XBB.1.16.9 | Spain | 0.949 | 0.935 | 0.964 | 1.000 | 4123.263 | 9315.2 |
| JA.1 | Spain | 0.946 | 0.935 | 0.956 | 1.001 | 2071.001 | 5598.9 |
| FU.1 | Spain | 0.944 | 0.933 | 0.955 | 1.001 | 2080.007 | 5416.7 |
| XBB.1.9 | Spain | 0.935 | 0.920 | 0.951 | 1.000 | 3791.110 | 8723.2 |
| XBB.1.16 | Spain | 0.930 | 0.924 | 0.936 | 1.003 | 610.683 | 1629.4 |
| XBB.2.3.2 | Spain | 0.929 | 0.919 | 0.940 | 1.001 | 1978.810 | 5019.6 |
| XBB.1.16.21 | Spain | 0.927 | 0.915 | 0.938 | 1.000 | 2332.198 | 6173.2 |
| XBB.1.16.1 | Spain | 0.924 | 0.916 | 0.932 | 1.001 | 1093.883 | 2955.5 |
| XBB.1.16.19 | Spain | 0.924 | 0.913 | 0.935 | 1.000 | 2328.623 | 6461.3 |
| FE.1.2 | Spain | 0.920 | 0.910 | 0.929 | 1.001 | 1649.337 | 4474.2 |
| XBB.2.3.3 | Spain | 0.916 | 0.905 | 0.927 | 1.002 | 1904.141 | 6021.5 |
| FE.1.1 | Spain | 0.912 | 0.901 | 0.923 | 1.001 | 1953.850 | 6181.3 |
| XBB.1.5.59 | Spain | 0.907 | 0.899 | 0.915 | 1.001 | 1032.917 | 3194.1 |
| XBB.2.3.14 | Spain | 0.903 | 0.891 | 0.915 | 1.000 | 2759.249 | 7050.5 |
| FY.6 | Spain | 0.903 | 0.889 | 0.917 | 1.000 | 3580.641 | 7655.5 |
| XBB.1.5.28 | Spain | 0.903 | 0.892 | 0.913 | 1.001 | 2063.777 | 5391.4 |
| XBB.2.3 | Spain | 0.899 | 0.893 | 0.905 | 1.003 | 604.509 | 1740.3 |
| EU.1.1 | Spain | 0.898 | 0.886 | 0.910 | 1.001 | 2223.916 | 5608.2 |
| XBB.1.16.2 | Spain | 0.897 | 0.886 | 0.908 | 1.001 | 2394.894 | 6935.9 |
| GB.1 | Spain | 0.897 | 0.884 | 0.909 | 1.000 | 2626.794 | 6258.7 |
| EU.1.1.1 | Spain | 0.896 | 0.882 | 0.911 | 1.000 | 3530.884 | 7409.5 |
| XBB.1.5.71 | Spain | 0.894 | 0.887 | 0.902 | 1.002 | 888.203 | 2928.0 |
| EG.14 | Spain | 0.894 | 0.880 | 0.906 | 1.001 | 3159.082 | 6463.0 |
| XBB.2.3.13 | Spain | 0.892 | 0.884 | 0.900 | 1.001 | 1121.797 | 3459.9 |
| FL.13 | Spain | 0.892 | 0.879 | 0.904 | 1.001 | 2904.163 | 6527.0 |
| FL.4 | Spain | 0.887 | 0.881 | 0.893 | 1.002 | 762.202 | 2149.6 |
| XBB.1.9.1 | Spain | 0.884 | 0.878 | 0.890 | 1.002 | 684.272 | 2071.7 |

| | | | | | | | |
|---|---|---|---|---|---|---|---|
| XBB.1.5.16 | Spain | 0.884 | 0.869 | 0.897 | 1.001 | 3468.149 | 6577.9 |
| EG.1.5 | Spain | 0.882 | 0.870 | 0.893 | 1.001 | 2325.130 | 6529.0 |
| FL.2 | Spain | 0.881 | 0.868 | 0.893 | 1.001 | 2467.962 | 5914.7 |
| BQ.1.1 | Spain | 0.880 | 0.866 | 0.893 | 1.001 | 3067.397 | 7905.3 |
| EG.1 | Spain | 0.879 | 0.871 | 0.886 | 1.002 | 930.927 | 2981.9 |
| XBB.1.5.4 | Spain | 0.876 | 0.862 | 0.890 | 1.000 | 3318.587 | 8020.5 |
| FL.1 | Spain | 0.875 | 0.861 | 0.889 | 1.001 | 3288.947 | 6671.7 |
| XBB.1.5.18 | Spain | 0.873 | 0.857 | 0.888 | 1.001 | 3828.278 | 8039.2 |
| XBB.1.5.37 | Spain | 0.871 | 0.864 | 0.877 | 1.002 | 753.260 | 2014.1 |
| EG.13 | Spain | 0.870 | 0.857 | 0.883 | 1.001 | 2749.987 | 7667.8 |
| HT.2 | Spain | 0.870 | 0.851 | 0.888 | 1.000 | 5162.995 | 10937.5 |
| GF.1 | Spain | 0.869 | 0.851 | 0.887 | 1.000 | 4950.329 | 8591.6 |
| XBB.1.9.2 | Spain | 0.867 | 0.860 | 0.874 | 1.002 | 818.926 | 2380.6 |
| EL.1 | Spain | 0.867 | 0.854 | 0.879 | 1.000 | 3024.317 | 7929.2 |
| XBB.1.5.20 | Spain | 0.866 | 0.848 | 0.883 | 1.000 | 5124.331 | 9070.8 |
| FL.3.2 | Spain | 0.863 | 0.856 | 0.871 | 1.002 | 980.825 | 2905.3 |
| XBB.1.5.67 | Spain | 0.863 | 0.848 | 0.878 | 1.001 | 3769.758 | 8247.0 |
| FL.5 | Spain | 0.862 | 0.851 | 0.873 | 1.001 | 2504.378 | 6741.1 |
| XBB.1.5 | Spain | 0.861 | 0.856 | 0.866 | 1.004 | 520.673 | 1365.6 |
| EG.1.3 | Spain | 0.859 | 0.844 | 0.873 | 1.000 | 3119.715 | 7887.8 |
| XBB.1.19.1 | Spain | 0.859 | 0.840 | 0.876 | 1.000 | 5765.932 | 10471.2 |
| XBB.1.5.65 | Spain | 0.857 | 0.837 | 0.876 | 1.000 | 5630.934 | 10593.9 |
| EG.1.6 | Spain | 0.856 | 0.841 | 0.871 | 1.000 | 3834.677 | 8538.0 |
| XBB.1.5.1 | Spain | 0.855 | 0.843 | 0.867 | 1.001 | 2753.409 | 7346.3 |
| FL.3 | Spain | 0.855 | 0.841 | 0.868 | 1.001 | 3550.104 | 8250.2 |
| XBB.1.17.1 | Spain | 0.854 | 0.840 | 0.868 | 1.001 | 3390.794 | 8328.2 |
| EG.1.2 | Spain | 0.850 | 0.829 | 0.869 | 1.001 | 5973.179 | 10449.2 |
| XBB.1.5.13 | Spain | 0.847 | 0.832 | 0.862 | 1.000 | 3637.820 | 7904.5 |
| XBB.1.5.12 | Spain | 0.844 | 0.823 | 0.864 | 1.000 | 7147.019 | 10955.7 |
| FG.2 | Spain | 0.844 | 0.828 | 0.859 | 1.000 | 4633.244 | 9087.6 |
| XBB.1.5.23 | Spain | 0.843 | 0.820 | 0.864 | 1.001 | 8345.020 | 10299.3 |
| FL.12 | Spain | 0.843 | 0.820 | 0.864 | 1.000 | 7177.196 | 11725.7 |
| CH.1.1 | Spain | 0.841 | 0.818 | 0.863 | 1.000 | 6967.314 | 10415.5 |
| XBB.2.4 | Spain | 0.839 | 0.826 | 0.851 | 1.001 | 2810.110 | 7322.5 |
| XBB.1.5.56 | Spain | 0.834 | 0.811 | 0.856 | 1.000 | 7887.374 | 11448.1 |
| FL.31 | Spain | 0.834 | 0.810 | 0.857 | 1.000 | 7755.363 | 11199.6 |
| XBB.1.5.77 | Spain | 0.832 | 0.816 | 0.848 | 1.000 | 4851.111 | 9626.1 |
| XBB.1.5.7 | Spain | 0.832 | 0.819 | 0.844 | 1.000 | 2994.563 | 6274.7 |
| XBB.1.5.5 | Spain | 0.827 | 0.804 | 0.848 | 1.000 | 7615.124 | 10759.6 |
| XBB.1.5.24 | Spain | 0.825 | 0.803 | 0.847 | 1.000 | 8807.286 | 11684.5 |
| XBB.1.5.66 | Spain | 0.816 | 0.789 | 0.842 | 1.000 | 9879.463 | 12089.8 |
| XBB.1.5.48 | Spain | 0.784 | 0.751 | 0.816 | 1.000 | 16915.282 | 11449.9 |
| FL.1.2 | Spain | 0.744 | 0.705 | 0.781 | 1.000 | 17379.887 | 11637.6 |

**Table S4. Primers used in this study**

| Primer name | Primer sequence (5'-to-3') | Purpose |
|---|---|---|
| Omicron universal Fw | cactatagggcgaattgggtaccatgtttgtgttcctggt | Preparation of S expression plasmid |
| BA.2 WT Rv | agctccaccgcggtggcggccgctcaggtgtagtgcagtttca | Preparation of S expression plasmid |
| pC-S_BA286_L455S_Fw | GACtacTGGtacagaagcttcaggaagagcAAG | Preparation of S expression plasmid |
| pC-S_BA286_L455S_Rv | CTTgctcttcctgaagcttctgtaCCAgtaGTC | Preparation of S expression plasmid |
| L455S_RBD_Fw | TTCTAAGCATAGTGGTAATTATGATTACTGGT ATAGATCGTTTAGGAAGTCTAAACTCAAACCT TTTGAGAG | Preparation of RBD expression plasmid |
| L455S_RBD_Rv | CATGGGAAAACATGTTGTTTACGGAG | Preparation of RBD expression plasmid |

### Supplementary figure

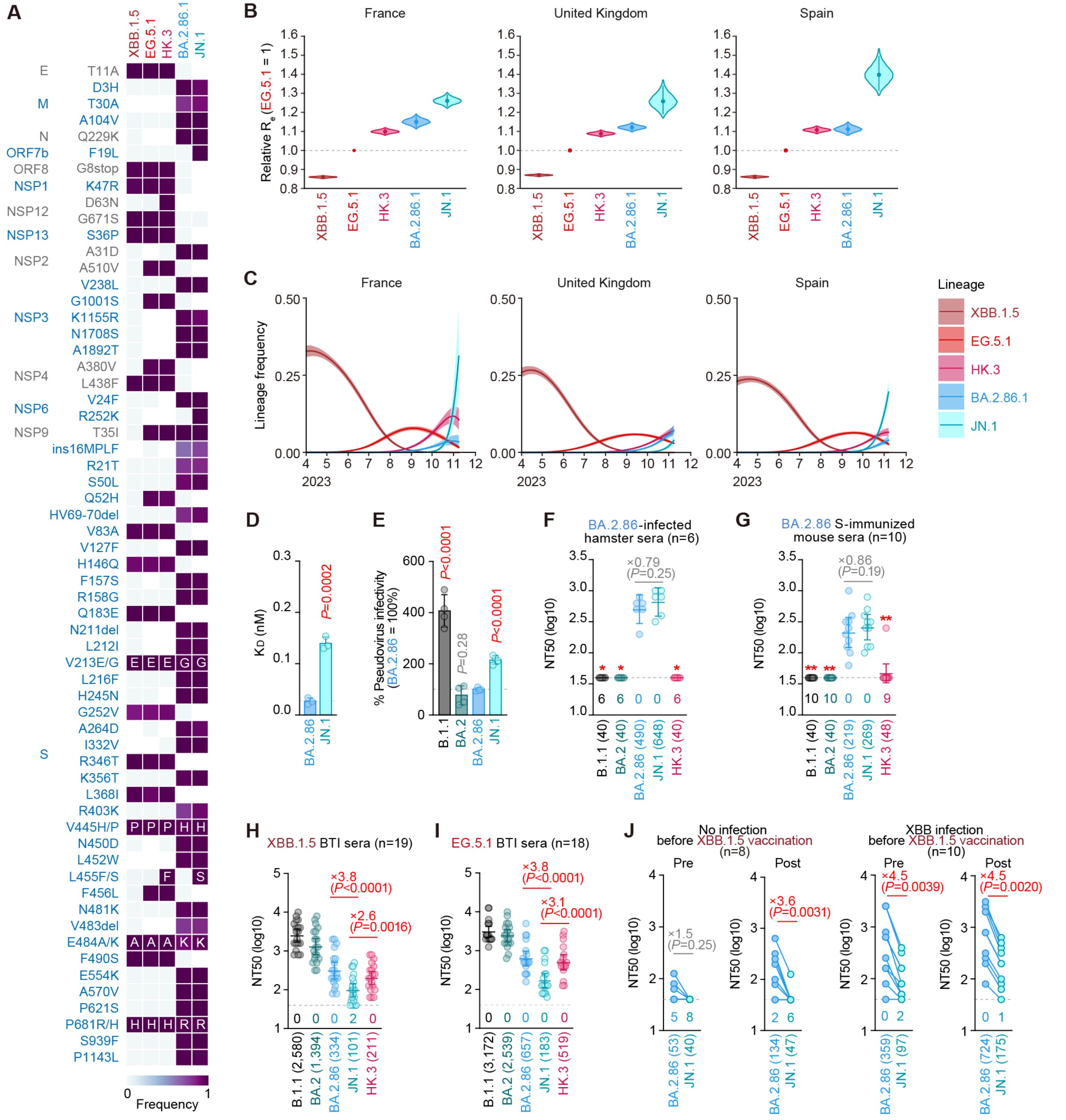


**Figure S1. Virological features of JN.1**

(**A**) Frequency of mutations in JN.1 and other lineages of interest. Only mutations with a frequency >0.5 in at least one but not all the representative lineages are shown.

(**B**) Estimated relative $R_e$ of the variants of interest in France, United Kingdom, and Spain. The relative Re of EG.5.1 is set to 1 (horizontal dashed line). Violin, posterior distribution; dot, posterior mean; line, 95% Bayesian confidence interval.

(**C**) Estimated epidemic dynamics of the variants of interest in France, United Kingdom, and Spain from April 1, 2023 to November 16, 2023. Countries are ordered according to the number of detected sequences of JN.1 from high to low. Line, posterior mean, ribbon, 95% Bayesian confidence interval.

(**D**) Yeast surface display affinity between the RBD of the BA.2.86 SARS-CoV-2 variant or BA.2.86 that contained the L455S mutation and mACE2 was measured by yeast surface display. The dissociation constant ($K_D$) value indicates the binding affinity of the RBD of the SARS-CoV-2 S protein to soluble ACE2 when expressed on yeast. Statistically significant differences versus BA.2.86 is determined by two-sided Student's *t* tests.

(**E**) Lentivirus-based pseudovirus assay. HOS-ACE2/TMPRSS2 cells were infected with pseudoviruses bearing each S protein of B.1.1 or BA.2 sublineages. The amount of input virus was normalized to the amount of HIV-1 p24 capsid protein. The percentage infectivity of B.1.1, BA.2 and JN.1 are compared to that of BA.2.86. The horizontal dash line indicates the mean value of the percentage infectivity of BA.2.86. Assays were performed in quadruplicate, and a representative result of four independent assays is shown. The presented data are expressed as the average ± SD. Each dot indicates the result of an individual replicate. Statistically significant differences versus BA.2.86 is determined by two-sided Student's *t* tests.

(**F**-**J**) Neutralization assay. Assays were performed with pseudoviruses harboring the S proteins of B.1.1, BA.2, BA.2.86, JN.1 and HK.3. The following sera were used: sera from six hamsters infected with BA.2.86 (**F**); sera from ten mice immunized with SARS-CoV-2 BA.2.86 S (**G**); convalescent sera from fully vaccinated individuals who had been infected with XBB.1.5 (eight 3-dose vaccinated donors, six 4-dose vaccinated donors, four 5-dose vaccinated donors and one 6-dose vaccinated donor. 19 donors in total) (**H**); and EG.5.1 (one 2-dose vaccinated donor, four 3-dose vaccinated donors, five 4-dose vaccinated donors, four 5-dose vaccinated donors and four 6-dose vaccinated donors. 18 donors in total) (**I**). Assays were also performed with pseudoviruses harboring the S proteins of BA.2.86 and JN.1. The following two sera were used: vaccinated sera from fully vaccinated individuals who had not been infected (8 donors) and vaccinated sera from fully vaccinated individuals who had been infected with XBB subvariants (after June, 2023) (10 donors). Sera were collected before vaccination ('Pre') and 3-4 weeks after XBB.1.5 monovalent vaccination ('Post') (**J**). Assays for each serum sample were performed in triplicate to determine the 50% neutralization titer ($NT_{50}$).

Each dot represents one $NT_{50}$ value, and the geometric mean and 95% confidence interval are shown. The number in parenthesis indicates the geometric mean of $NT_{50}$ values. The horizontal dash line indicates the detection limit (40-fold) and the number of samples with neutralization titer under the limit are shown below the dash line. In **F**-**J**, statistically significant differences versus JN.1 were determined by two-sided Wilcoxon signed-rank tests, and p values are indicated in parentheses. The fold changes of $NT_{50}$ from that of JN.1 are indicated with "X". In **F** and **G**, *, $p<0.05$; **, $p<0.01$ versus JN.1.

**Consortia**

**The Genotype to Phenotype Japan (G2P-Japan) Consortium**


**The Institute of Medical Science, The University of Tokyo, Japan**
Naoko Misawa, Ziyi Guo, Jarel Elgin M. Tolentino, Shigeru Fujita, Lin Pan, Mai Suganami, Mika Chiba, Ryo Yoshimura, Kyoko Yasuda, Keiko Iida, Naomi Ohsumi, Adam P. Strange, Shiho Tanaka, Kaoru Usui, Wilaiporn Saikruang, Kaho Okumura

**Hokkaido University, Japan**
Takasuke Fukuhara, Tomokazu Tamura, Rigel Suzuki, Saori Suzuki, Hayato Ito, Keita Matsuno, Hirofumi Sawa, Naganori Nao, Shinya Tanaka, Masumi Tsuda, Lei Wang, Yoshikata Oda, Zannatul Ferdous, Kenji Shishido, Keita Mizuma, Isshu Kojima, Jingshu Li, Tomoya Tsubo, Shuhei Tsujino

**Tokai University, Japan**
So Nakagawa

**Kyoto University, Japan**
Kotaro Shirakawa, Akifumi Takaori-Kondo, Kayoko Nagata, Ryosuke Nomura, Yoshihito Horisawa, Yusuke Tashiro, Yugo Kawai, Kazuo Takayama, Rina Hashimoto, Sayaka Deguchi, Yukio Watanabe, Ayaka Sakamoto, Naoko Yasuhara, Takao Hashiguchi, Tateki Suzuki, Kanako Kimura, Jiei Sasaki, Yukari Nakajima, Hisano Yajima, Yoshitaka Nakata, Hiroki Futatsusako

**Hiroshima University, Japan**
Takashi Irie, Ryoko Kawabata

**Kyushu University, Japan**
Kaori Tabata

**Kumamoto University, Japan**
Terumasa Ikeda, Hesham Nasser, Ryo Shimizu, MST Monira Begum, Michael Jonathan, Yuka Mugita, Otowa Takahashi, Kimiko Ichihara, Takamasa Ueno, Chihiro Motozono, Mako Toyoda, Sharee Leong

**University of Miyazaki, Japan**
Akatsuki Saito, Maya Shofa, Yuki Shibatani, Tomoko Nishiuchi

**Tokyo Metropolitan Institute of Public Health, Japan**
Kazuhisa Yoshimura, Kenji Sadamasu, Mami Nagashima, Hiroyuki Asakura, Isao Yoshida

**Charles University, Czechia**
Prokopios Andrikopoulos, Miguel Padilla-Blanco, Aditi Konar

**Acknowledgments**

We would like to thank all members of The Genotype to Phenotype Japan (G2P-Japan) Consortium. We thank Dr. Kenzo Tokunaga (National Institute of Infectious Diseases, Japan) for sharing materials and Dr. Keita Matsuno (Hokkaido University, Japan) for sharing hamster sera. We gratefully acknowledge the numerous laboratories worldwide that have provided sequence data and metadata to GISAID. A full list of originating and submitting laboratories for the sequences used in our analysis can be found at https://www.gisaid.org using the EPI-SET-ID: EPI_SET_231110rt and EPI_SET_231111fo.
